# Cerebrospinal fluid inflammome analysis identifies host- and pathogen-specific inflammatory profiles and signaling pathways in meningitis and predicts clinical outcome

**DOI:** 10.1101/2025.03.27.645675

**Authors:** Thorsten Lenhard, Marie-Therese Herkel, Christine S. Falk, Viktor Balzer, Gert Fricker, Corinna Schranz, Uta Meyding-Lamadé

**Affiliations:** Neuroinfectious Diseases Group, UniversityHospital, Heidelberg, Germany; Department of Neurology, UniversityHospital, Heidelberg, Germany; Institute of Transplant Immunology, Hannover Medical School, Hannover, Germany; Institute of Pharmacy and Molecular Biotechnology, University of Heidelberg, Germany; Department of Neurology, Krankenhaus Nordwest, Frankfurt, Germany; German Center for Infectious Diseases (DZIF, TTU-IICH), Braunschweig, Germany

## Abstract

Increasing evidence indicate that individual host immune response to pathogens may be as important as virulence factors in determining the severity of infection. However, most available data on pathogen-host interactions are based on in vitro findings, neglecting the genetic complexity of the immune response. We conducted a comparative observational study of bacterial and viral meningitis to identify pathogen-specific and host-dependent inflammatory pathways that may predict the severity of infection by analysing 54 inflammatory factors using plexing technology. Infection severity and neurological disability were assessed using the modified RANKIN scale. To identify pathogen-specific and host-dependent pathways, plex data were entered into EMBiology®™ software and gene set enrichment analysis (GSEA) was performed. We also investigated pathogen- and host-dependent regulatory effects on blood-brain barrier function using human iPSC-derived endothelial cells as a model. BBB function was characterised at the level of transendothelial electrical resistance (TEER), endothelial apoptosis and drug transporter activity. We identified 36 factors that are highly differentially regulated in either bacterial or viral meningoencephalitis compared to healthy subjects, 15 of which have not been previously described. GSEA identified previously unknown pathogen-specific pathways including neurotrophin, NOTCH, immune tolerance, antiviral defence and tight junction signalling. As a key finding, we identified 15 host-dependent factors that correlated with the grade of disability and with blood-CSF barrier dysfunction. GSEA revealed stronger responses of neuroprotective (AKT/ERK), anti-apoptotic (Bcl-2/p53) and anti-inflammatory (JAK1/STAT1) pathways in patients with favourable course, whereas patients with severe infection show stronger pro-inflammatory STAT-activation, NKC/CTL-responses and NFκ-B-associated neurotoxicity. BBB analysis supports these findings with dysregulation of TEER, drug transporter activity and induction of endothelial cell death in patients with severe infection. Our data highlight the need for future approaches to personalised medicine that go beyond traditional anti-infective therapy to target the individual immune response.

**Author Summery:** Infections often cause personal suffering, but also high economic costs. Brain infections such as bacterial or viral meningitis are particularly serious and carry a high risk of death or permanent disability. At the same time, resistance to anti-infective drugs is increasing and fewer new antibiotics are being developed. As a result, there is a great need for translational research to develop new treatments beyond conventional anti-infectives. Due to its strict compartmentalisation, the brain is well suited to identify specific pathogen-dependent, but also host (patient) dependent inflammatory patterns in infections. Using a cohort of patients with a defined group of pathogen-specific meningitis, we investigated the inflammatory patterns in the cerebrospinal fluid and were able to describe new specific pathogen-dependent and new host-dependent immune responses and identify associated signalling pathways. We were also able to show how and at what cellular and molecular level the blood-brain barrier is disrupted by infection. The degree of damage to the blood-brain barrier is an important aspect and has major implications for the course and prognosis of brain infections. Once the pathogen- and host-specific aspects of the immune response are understood, new personalised therapeutic approaches can be developed that directly target the regulation of the individual immune response.

## Introduction

Global surveillance of acute meningoencephalitis has reported an increase in incidence over the past two decades [1]. In immunocompetent individuals, the majority of cases of acute meningoencephalitis are caused by bacteria (incidence: 3/100,000) or viruses (7/100,000) [2–4]. Worldwide incidence varies widely geographically and is strongly associated with poverty. For example, in meningitis belt countries, incidence reaches 1000/100,000 cases and is almost exclusively caused by *Neisseria meningitidis* [1]. The incidence of viral encephalitis also varies geographically, mainly due to vector-borne infections such as Japanese encephalitis in Southeast Asia or tick-borne encephalitis in Europe [5, 6]. Either way, acute meningoencephalitis causes a significant number of severe infections resulting in chronic disability or even death. Pathogen-derived virulence factors determine the severity of infection and clinical outcome. Numerous examples of pathogenicity factors have been described in bacteria (e.g. M protein in *Streptococcus species*) and viruses (e.g. HSV-1 ICP47 as an inhibitor of MHC I, HSV-1 ICP27 as an inhibitor of host RNA processing or poliovirus 2B/3A as an inhibitor of host ER-to-Golgi protein traffic), to name but a few [7–10]. In addition, in some bacteria, epigenetic mobile elements such as plasmid- and phage-encoded factors are important determinants of virulence (e.g. diphteria toxin in *Corynebacterium diphteriae*) [11]. Thus, variations in pathogen strains and host immunological competence are thought to determine the course and severity of infection. However, less is known about the host (patient) side and the factors that determine the clinical course and outcome of infections. For example, at the single gene level, polymorphisms in the CCR5 gene are associated with severe West Nile encephalitis but also confer resistance to HIV [12]. Mutations in the IFNγ-signaling pathway gene Unc-93B cause much higher susceptibility to HSV-1 [13]. However, the complex host-pathogen interactions in vivo are poorly understood and there is a great need for translational research to better understand these interactions. Particularly from the host’s perspective, a balanced immune response will ensure the best possible outcome by controlling pathogen proliferation and enabling pathogen clearance as effectively as possible. However, terminating the immune response in an appropriate manner is also important to prevent secondary damage caused by an excessive immune response [14, 15].

The brain is immunologically privileged because the blood-brain barrier (BBB) controls the transport of immunoglobulins and the transmigration of immune cells and also serves as a barrier to pathogens [16]. At the molecular level, drug transporters expressed in the brain endothelium protect the brain from endogenous and xenogenous toxins [17]. The BBB is a multicellular and molecular structure defined by the neurovascular unit (NVU), which includes endothelial cells, pericytes, astrocyte endfeet and hemisynapses, but also extracellular molecular structures such as the basal membrane, extracellular matrix and apical glycocalyx [18]. Brain infections can cause indirect and direct damage to the NVU from both the blood and the subendothelial brain tissue sides, leading to increased brain dysfunction [19]. In addition, systemic infections have remote effects on BBB function, such as sepsis-associated encephalopathy (SAE) or cerebral malaria, both of which are associated with increased morbidity and mortality [20–22].

Because of its strict compartmentalisation, the brain is an ideal organ to study both pathogen-specific and host-specific parts of the immune response. Accordingly, we performed a comprehensive analysis of brain inflammation in a cohort of immunocompetent patients with bacterial meningoencephalitis (BME) and viral meningoencephalitis (VME) and compared the responses with those of healthy controls using multiplex bead array technology. We also performed deep network analysis using gene set enrichment analysis (GSEA) and downstream pathway analysis to identify specific inflammatory pathways that are embedded in the specific inflammatory response to a given pathogen and to identify specific host (patient) related inflammatory patterns and their associated pathways that may be associated with the severity of infection [23, 24]. To broaden the focus to a critical site of infection, we further investigated the dysregulation of the BBB as a critical structure normally responsible for maintaining intact brain homeostasis. To this end, we investigated a human in vitro BBB model based on differentiated human brain endothelial cells derived from inducible pluripotent stem cells (iPSCs). We analysed the regulation of cellular and molecular properties of the BBB such as transendothelial electrical resistance (TEER), drug transporter activities and inducible endothelial cell death.

## Results

### Infectious cohorts

Causative pathogens of BME cohort: *Streptococcus pneumonia* (9), other *Streptococcus spec.* (5), *Staphylococcus aureus* (5), *Neisseria meningitides* (4), and other species (4). VME cohort: 12 *Herpesviridae*-derived infections composed of VZV (6), HSV-1 (4), and others (2) and 11 RNA viruses composed of tick-borne encephalitis virus (TBEV) (9) and Enteroviruses (2). All patients suffered from at least one focal neurological symptom, from disturbance of consciousness or from focal epileptic seizure or combinations thereof to fulfil criteria for brain parenchyma involvement (=encephalitis) [25]. The subgroups (VME, BME) were comparable in age and gender distribution, but differed in CSF cell count and albumin quotient. There was a non-significant trend in the severity of infection in the pathogen groups as measured by the modified Rankin Scale (mRS) [26]. **Table 1** summarises the baseline characteristics of the different cohorts.

**Table 1.** Baseline characteristics of the viral and bacterial infection cohorts and the healthy control group.

|  | Age |  | Sex |  | mRS |  | QAlb |  | CSF<br>cell number |  |
| --- | --- | --- | --- | --- | --- | --- | --- | --- | --- | --- |
|  | median | P-value | male [%] |  | median | P-value | mean | P-value | median | P-value |
| **Infectious cohorts**<br>n=50 |  |  |  |  |  |  |  |  |  |  |
| VME<br>n=23 | 57 | ns<sup>§</sup> | 55 |  | 4 | 0.005<sup>§</sup> | 18,0 | < 0.001<sup>§</sup> | 259 | < 0.001<sup>§</sup> |
| BME<br>n=27 | 54 | ns<sup>§,#</sup> | 56 |  | 5 | 0.003<sup>§</sup><br>0.051<sup>#</sup> | 126,3 | < 0.001<sup>§</sup><br>< 0.001<sup>#</sup> | 3306 | < 0.001<sup>§</sup><br>0.003<sup>#</sup> |
| **Control cohort**<br>n=32 | 51 |  | 59 |  | 0 |  | 6,2 |  | 1 |  |

mRS, modified RANKIN scale; QAlb, CSF/serum albumin quotient; CSF, cerebrospinal fluid; VME, viral meningoencephalitis; BME, bacterial meningoencephalitis; ns , not significant; §, compared to control; #, BME compared to VME.

### Pathogen-dependent inflammation in BME and VME

**S1 Table** shows the 54 investigated soluble inflammatory factors. None of the factors correlated with age or were significantly differentially expressed between the sexes (see correlation heat map, **S1 Fig.**). Thirty-six factors (67%) of the 54 investigated factors were significantly and differentially up- or downregulated in either BME, VME or both. Average regulation across all factors compared to control: 100’fold in BME with a range of 2.1-1322’fold, ǀPǀ=0.045-0.0000001 and 197’fold in VME with a range of 1.39-4947’fold, ǀPǀ=0.048-0.000004. Of these, 40% correlated either positively or negatively with the level of disability as measured by the mRS (**S1 Fig.**). In addition, there was a moderate correlation between QAlb and the level of disability measured as mRS and a weak correlation between pleocytosis and mRS (see correlation heat map, **S1 Fig.**) [26]. Furthermore, there was a strong inverse correlation between CSF glucose concentration and mRS, but only in the BME group. **Table 2** shows the inflammatory factors that were exclusively differentially regulated in either BME or VME, of which 58% additionally correlated negatively or positively with grade of disability mRS (see correlation heat map, **S1 Fig.** and **S2 Table**). Note that the identified regulated factors reflect exclusively intrathecal expression, as passive diffusion from outside the brain across the disrupted blood-brain barrier was excluded by factoring in QAlb (for calculation of the intrathecal factor proportion, see Materials and Methods section) [27–29].

**Table 2.**
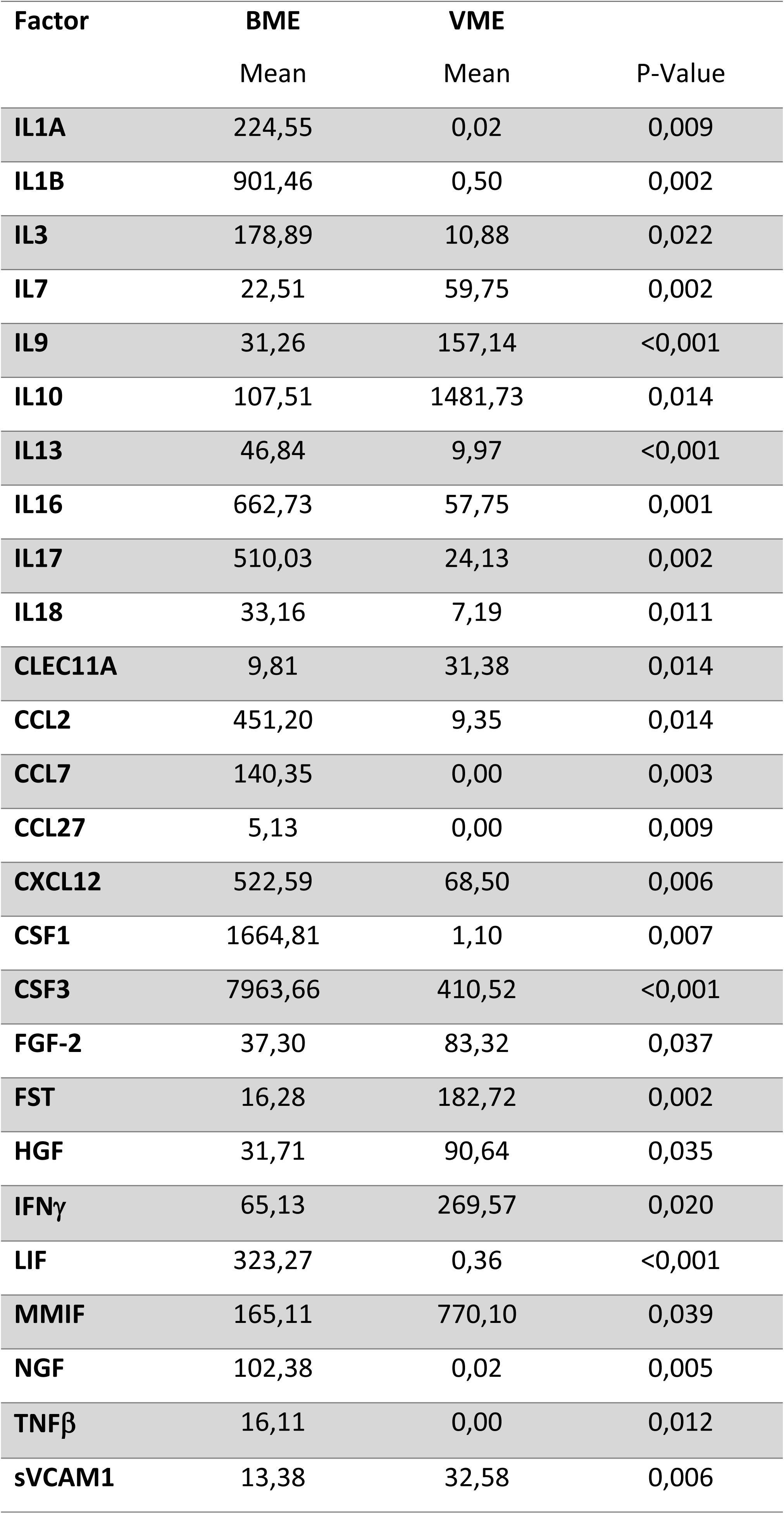
Expression of inflammatory factors in the cerebrospinal fluid in VME and BME.

| Factor | BME | VME | P-Value |
| --- | --- | --- | --- |
|  | Mean | Mean |  |
| **IL1A** | 224,55 | 0,02 | 0,009 |
| **IL1B** | 901,46 | 0,50 | 0,002 |
| **IL3** | 178,89 | 10,88 | 0,022 |
| **IL7** | 22,51 | 59,75 | 0,002 |
| **IL9** | 31,26 | 157,14 | <0,001 |
| **IL10** | 107,51 | 1481,73 | 0,014 |
| **IL13** | 46,84 | 9,97 | <0,001 |
| **IL16** | 662,73 | 57,75 | 0,001 |
| **IL17** | 510,03 | 24,13 | 0,002 |
| **IL18** | 33,16 | 7,19 | 0,011 |
| **CLEC11A** | 9,81 | 31,38 | 0,014 |
| **CCL2** | 451,20 | 9,35 | 0,014 |
| **CCL7** | 140,35 | 0,00 | 0,003 |
| **CCL27** | 5,13 | 0,00 | 0,009 |
| **CXCL12** | 522,59 | 68,50 | 0,006 |
| **CSF1** | 1664,81 | 1,10 | 0,007 |
| **CSF3** | 7963,66 | 410,52 | <0,001 |
| **FGF-2** | 37,30 | 83,32 | 0,037 |
| **FST** | 16,28 | 182,72 | 0,002 |
| **HGF** | 31,71 | 90,64 | 0,035 |
| **IFNγ** | 65,13 | 269,57 | 0,020 |
| **LIF** | 323,27 | 0,36 | <0,001 |
| **MMIF** | 165,11 | 770,10 | 0,039 |
| **NGF** | 102,38 | 0,02 | 0,005 |
| **TNFβ** | 16,11 | 0,00 | 0,012 |
| **sVCAM1** | 13,38 | 32,58 | 0,006 |

Mean values are calculated as specific intrathecal concentrations. Raw results for each factor in CSF and serum were originally reported in pg/ml. P values are presented as two-tailed t-tests. Factor names are given as gene symbols according to the HUGO approved human gene nomenclature https://www.genenames.org/.

### Pathogen-specific down-regulation of inflammatory factors and associated pathways

Having shown that a considerable number of inflammatory factors were differentially regulated in VME and BME, we were next interested in the patterns of downregulated factors in VME and in BME compared to levels in healthy controls. The rationale behind this approach was to identify those putative factors that might maintain physiological conditions in the brain, such as trophic support of neurons, but are downregulated in meningoencephalitis and thus fail to maintain neuronal homeostasis. In BME, a cluster of seven factors were found at significantly lower concentrations compared to healthy controls (**Fig 1)**. Gene set enrichment analysis (GSEA) identified five specific pathways associated with these identified factors in BME (**Fig 1B**), which can be classified as: cluster 1, inhibition and activation of apoptosis; cluster 2, neuroprotective growth factor signaling and associated downstream regulation of transcription factors; cluster 3, Notch signaling; cluster 4, immune tolerance and T-cell attraction; and cluster 5, matrix metalloproteinase activity. Note, since the here identified factors in BME were down-regulated compared to controls, likewise the associated pathways are supposed to be less active in BME.

**Fig 1.**
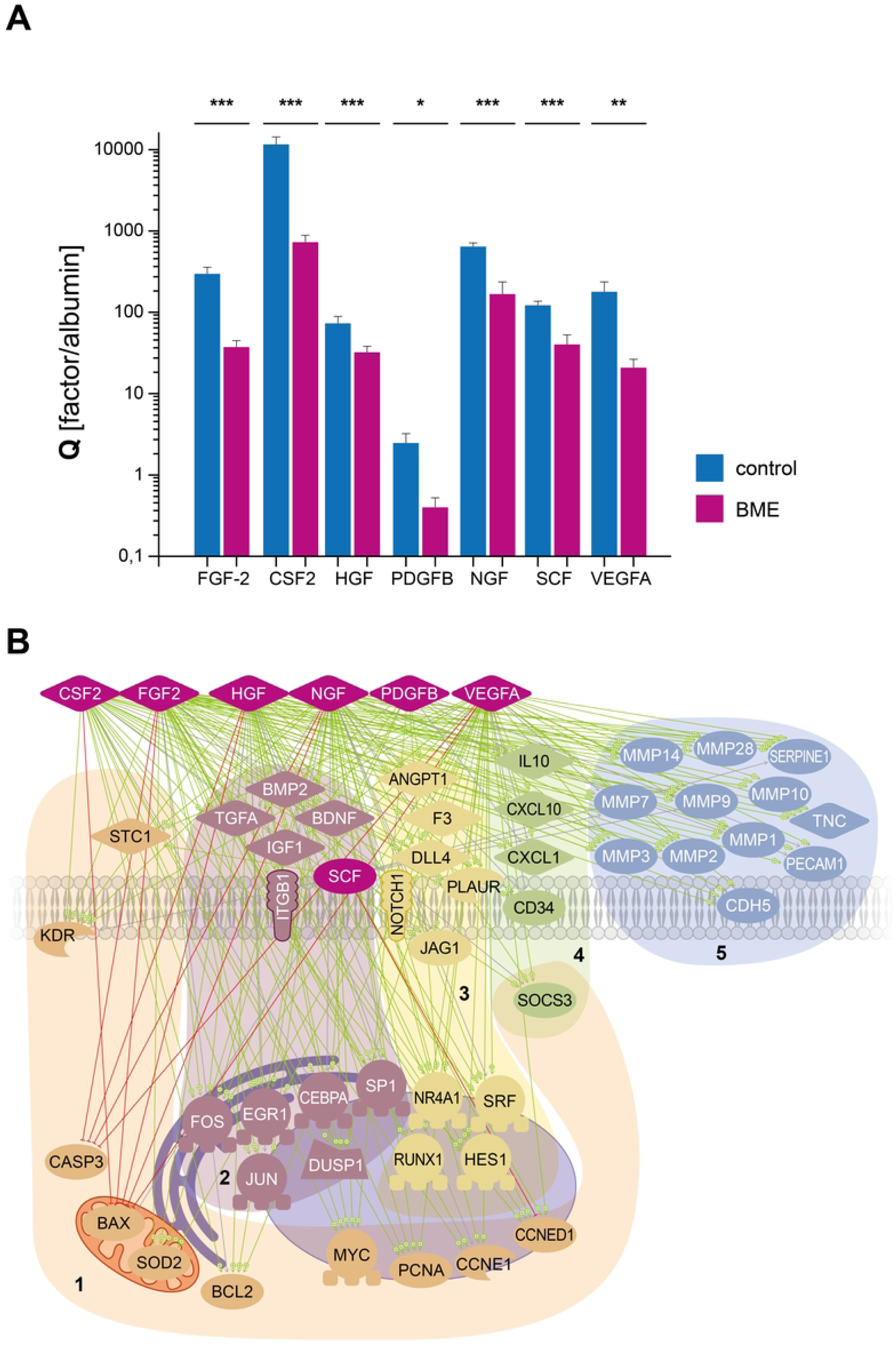
Downregulated inflammatory factors in bacterial meningoencephalitis and associated signaling pathways. Multiplex bead array analysis of intrathecally expressed inflammatory factors (A) and deduced signaling pathways (B) in bacterial menigoencephalitis (BME). In (A) factors are shown as specific quotient Q calculated from cerebrospinal fluid/serum factor concentration divided by albumin quotient to represent factors expressed exclusively intrathecally in BME compared to healthy controls. Note that the ordinate is logarithmically scaled. For detailed Q calculation and P-value allocation, see method section. Legend: FGF-2, basic fibroblast growth factor; CSF, colony stimulating factors; HGF, hepatocyte growth factor; PDGFB, platelet-derived growth factor B; NGF, nerve growth factor; SCF, stem cell factor; VEGFA, vascular endothelial growth factor A. In (B), gene set enrichment analysis (GSEA) enriched for the identified differentially regulated entities in BME is shown. The assignment of factors and targets to cellular and subcellular structures (cell membrane, mitochondria, ER, nucleus) is shown schematically. The downstream targets are grouped into specific clusters: 1, apoptotic signals; 2, neurotrophic factor and associated transcription factor signals; 3, Notch signals; 4, immunotolerance signals; 5, matrix metalloproteinase signals.Protein symbols: 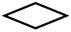, growth factors, interleukins, cytokines, chemokines; 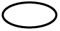, regulatory proteins (e.g. proteases); 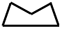, regulatory phosphatases; 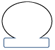, transcription factors; 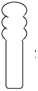, receptors. All molecules are shown as gene symbols, for full name, see http://www.genenames.org.

In VME, a cluster of five factors was significantly downregulated compared to healthy controls (**Fig 2**). GSEA revealed six differentially targeted pathways compared to controls (**Fig 2B**). Three pathways are basically similar to those found in BME (cluster 1, 2, 5), but three additional pathways were different: cluster 4, antiviral defence factors; cluster 5, chemokine signaling; and cluster 6, tight junction regulation. As already pointed for BME, these associated pathways are supposed to be likewise less active in VME.

**Fig 2.**
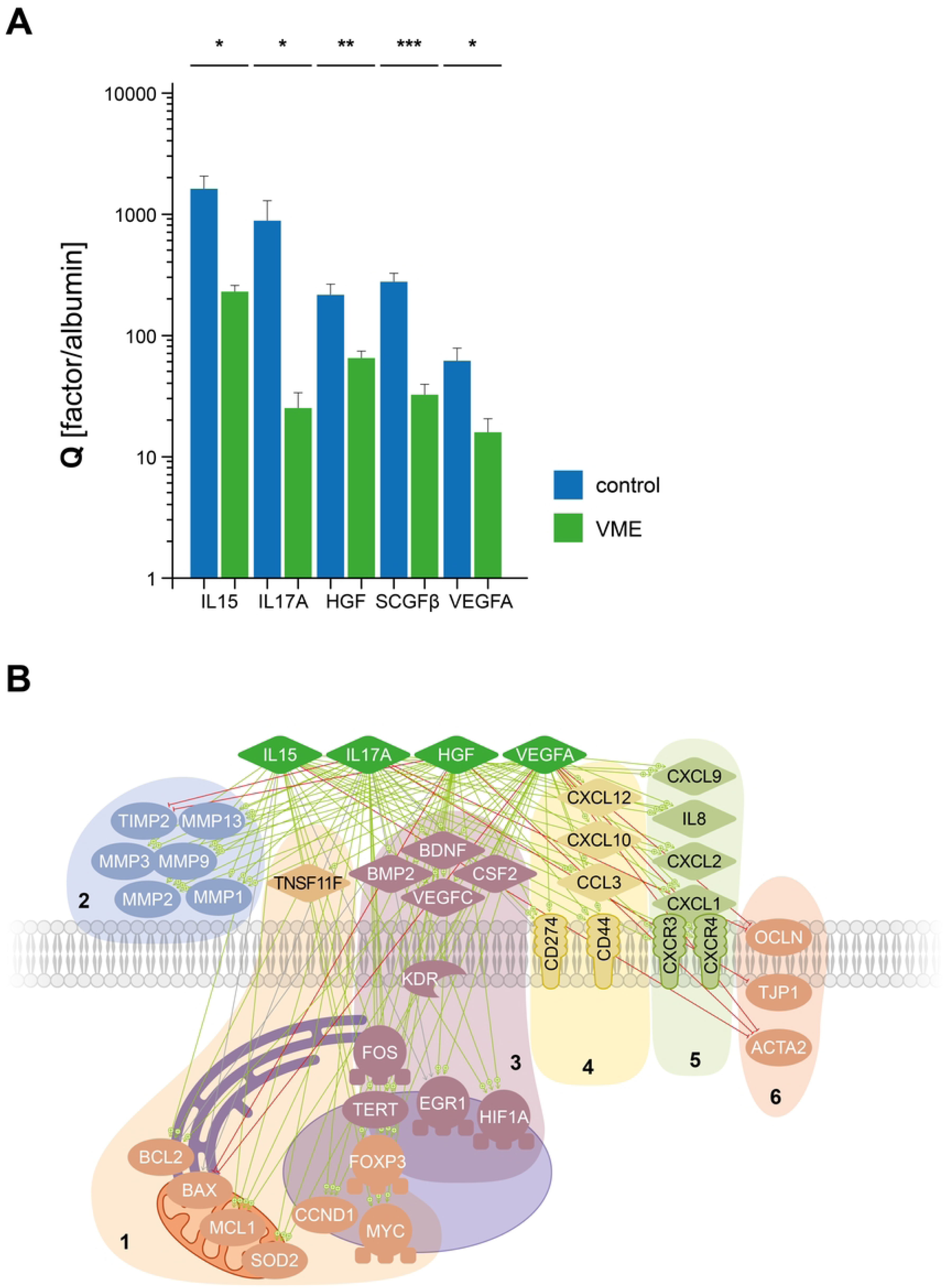
Downregulated inflammatory factors in viral meningoencephalitis and associated signaling pathways. Multiplex bead array analysis of intrathecally expressed inflammatory factors (A) and deduced signaling pathways (B) in viral menigoencephalitis (VME). As in Fig 1, the assignment of factors and targets to cellular and subcellular structures (cell membrane, mitochondria, ER, nucleus) is shown schematically. The downstream targets a grouped into specific clusters: for protein symbols and cluster 1 - 3, compare figure 1; 4, antiviral defense signals; 5, chemokine signalling; 6, tight junction signals. Legend: IL, interleukin; SCGFβ, stem cell growth factor-beta, for detailed description, see Fig 1. All molecules are shown as gene symbols, for full name, see http://www.genenames.org.

### Differentially regulated inflammation and associated pathways as a function of patient’s disability

In particular, we were interested in the regulation of inflammatory factors as a function of disability in patients, to identify patterns that might be associated with outcome regardless of the pathogen. We therefore dichotomised the cohort, irrespective of whether a patient had VME or BME, by median disability (mRS) into a mild to moderate (“favourable”) subgroup (mRS ≤ 4, median = 3, range 2-4) and a severely affected (“unfavourable”) subgroup (mRS > 4, median = 5, range 5-6). To reduce pathogen-specific effects of BME and VME, respectively, and to extract host-specific patterns, we defined mRS-matched pairs of VME and BME, i.e. each VME with a given mRS score was paired with a BME case with the same mRS score. This allowed us to include 23 pairs in the final subgroup analysis. We then performed subgroup analyses for differentially regulated inflammatory factors. **Table 3** summarize the significantly regulated factors of both the favourable (mRS ≤ 4) and unfavourable (mRS > 4) subgroups. **Figure 3** shows a correlation heatmap (spearman-rho) of these identified factors. Correlation analysis of each factor showed weak to strong positive or negative correlation with both subgroups (favourable *versus* unfavourable), and each factor also correlated with the grade of mRS itself. In addition, 53% of the factors (IL12B, IL15, IFNα2, ICAM1, VCAM1, CSF3, SCF and CLEC11A) also correlated weakly to moderately with the degree of blood-CSF barrier dysfunction as measured by QAlb. Unfortunately, pathway analysis of the subgroups using the EMBiology^®^™ software was not possible due to the small total number of patients included in the subgroup analysis. Therefore, a structured Boolean search strategy was performed in PubMed to identify putative associated pathways. Further details of the search strategy and results can be found in the **S1 Methods Appendix** and **S3 Table**. **Figure 4** summarises the putative pathways involved in patients with a favourable (mRS ≤ 4) and unfavourable (mRS > 4) course of infection. In the favourable subgroup, pathways involved in immune response termination, neuroprotection and regulation of apoptosis were identified (**Fig 4A**), whereas in the unfavourable subgroup, pathways involved in INFγ signaling, neurotoxic pathways and cytotoxic T cell/NK cell response pathways were identified (**Fig 4B**).

**Fig 3.**
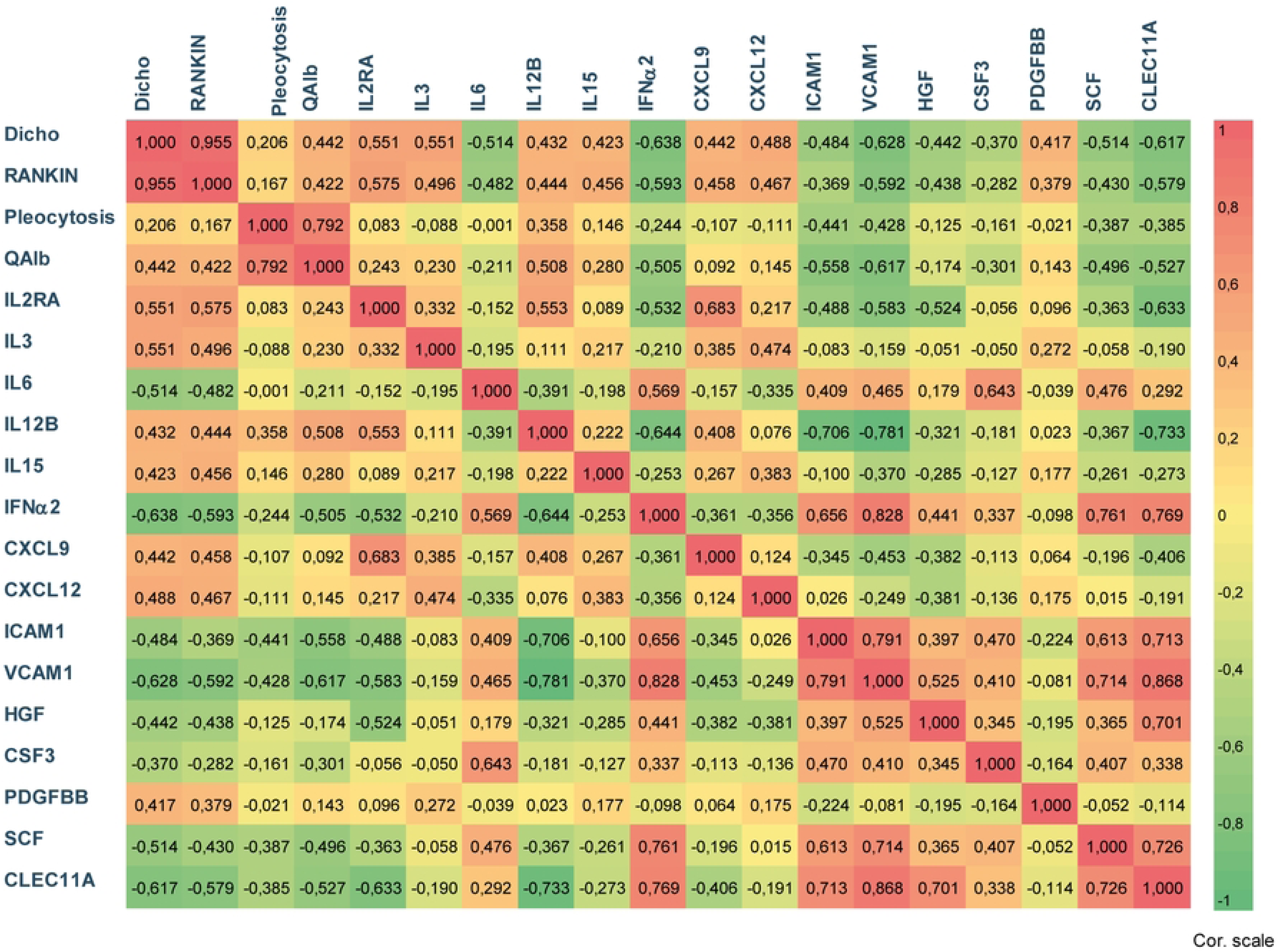
Correlation heatmap of regulated inflammatory factors as a function of host (patient) disability. Shown is a correlation heatmap of a Spearman rho correlation. The correlation coefficient (ρ) is shown for each pair of variables. ρ-values between +/-0.395 - +/-0.5 indicate a weak correlation and show P-values < 0.05 (2-sided), ρ-values between +/-0.5 - +/-0.868 indicate a moderate to strong correlation and show P-values < 0.01 (2-sided). Dicho is defined as favourable infection course with mRS ≤ 4 or unfavourable course with mRS > 4. Note, that variables correlating with unfavourable course are coloured in shades of red and variables correlating with favourable course are coloured in shades of green (only first two column and first two lines, respectively). RANKIN, modified RANKIN scale (mRS); QAlb, CSF albumin quotient; Cor. scale, correlation coefficient scale (ρ); inflammatory factors and growth factors, all molecules are shown as gene symbols, for details see also http://www.genenames.org.

**Fig 4.**
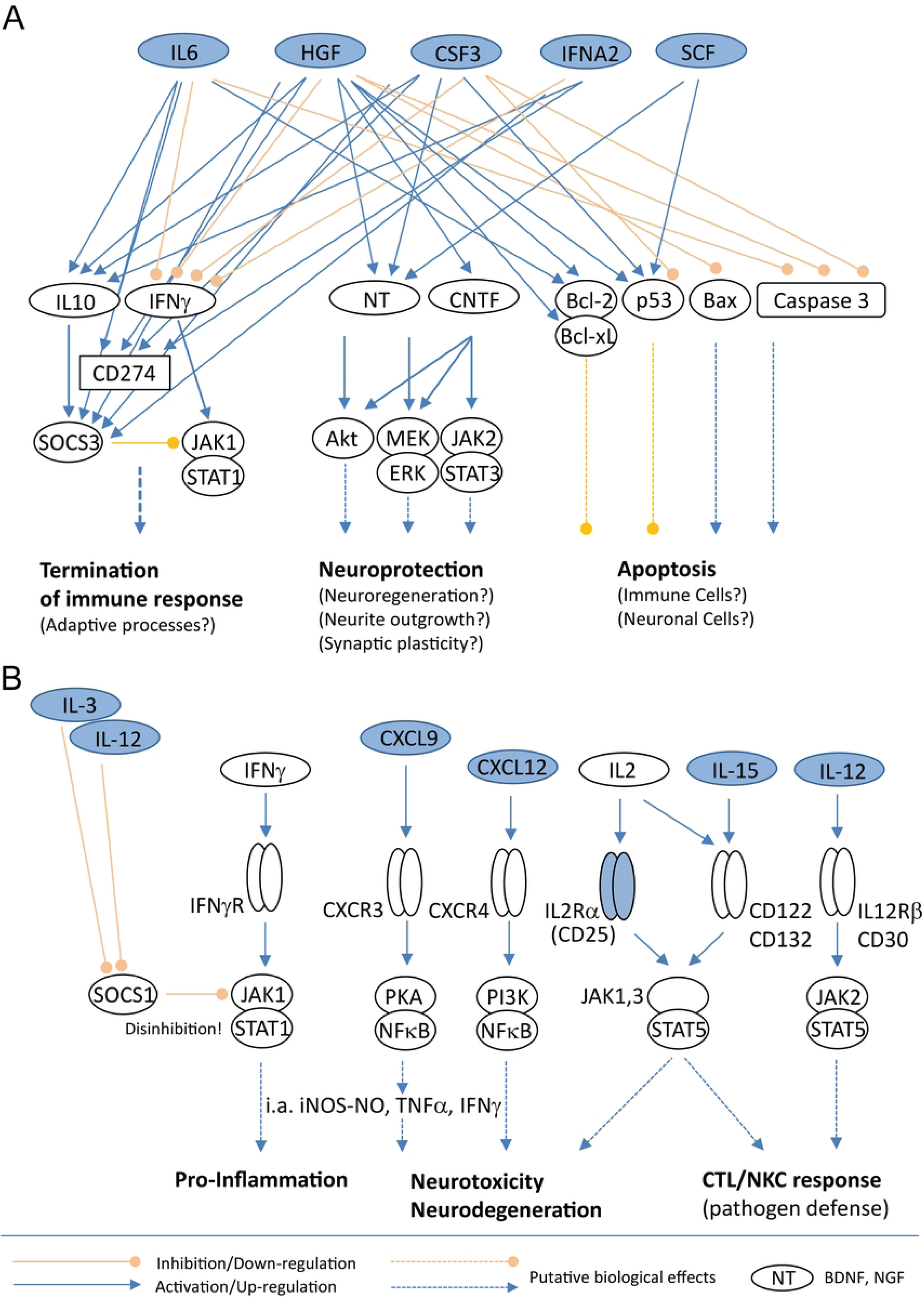
Host (patient) dependent signaling pathways and associated biological functions in patients with meningoencephalitis. Shown are identified pathways involved in either a favourable (mRS ≤ 4) (A) or unfavourable (mRS > 4) (B) course of infection based on the Boolean search strategy (for details, see S1 Methods Appendix and S3 Table). Factors shown in blue are those measured to be regulated in our subgroups. Legend: IL, interleukin; HGF, hepatocyte growth factor; CSF3, colony stimulating factor 3 (G-CSF); IFNA2, interferon-alpha2; SCF, stem cell factor; CXCL, chemokine (C-X-C motif) ligand; IFNγ, interferon-gamma; CD, cluster of differentiation; CNTF, ciliary neurotrophic factor; Bcl, B-cell lymphoma; Bax, Bcl-2-associated X protein; SOCS, suppressor of cytokine signaling; JAK-STAT, Janus kinase (JAK)-signal transducer and activator of transcription (STAT) pathway; Akt, protein kinase B; MEK-ERK, mitogen-activated protein kinase kinase (MEK)/extracellular signal-regulated kinase (ERK) pathway; NFκ-B, nuclear factor ’kappa-light-chain-enhancer’ of activated B cells; PKA, protein kinase A; R, abbreviation for receptor; all molecules are shown as gene symbols, for details see also http://www.genenames.org.

**Table 3.** Expression of CSF factors in patients with favourable and unfavourable course of infection.

| Factor/subgroup | RS $\leq 4$ | RS $> 4$ | P-Value |
| --- | --- | --- | --- |
|  | Mean | Mean |  |
| **sIL2RA** | 59,65 | **164,13** | 0,007 |
| **IL3** | 43,46 | **138,76** | 0,005 |
| **IL6** | **33863,73** | 4395,48 | 0,038 |
| **IL12B** | 278,39 | **863,12** | 0,038 |
| **IL15** | 4,02 | **12,86** | 0,018 |
| **IFNα2** | **384,74** | 54,23 | 0,037 |
| **CXCL9** | 1083,04 | **3857,83** | 0,017 |
| **CXCL12** | 6,25 | **32,36** | 0,035 |
| **sICAM1** | **12,89** | 5,86 | 0,045 |
| **sVCAM1** | **36,41** | 10,48 | 0,002 |
| **HGF** | **77,56** | 34,47 | 0,029 |
| **CSF3** | **2086,46** | 427,83 | 0,039 |
| **PDGFB** | 0,38 | **1,13** | 0,035 |
| **KL (SCF)** | **90,75** | 27,80 | 0,046 |
| **CLEC11A (SCGFβ)** | **42,19** | 9,20 | 0,032 |

Mean values are calculated as specific intrathecal concentrations. Raw results for each factor in CSF and serum were originally reported in pg/ml. P values are presented as two-tailed t-tests. Factor names are given as gene symbols according to the HUGO approved human gene nomenclature https://www.genenames.org/.

### Dysregulation of blood-brain-barrier function

Differentiated human pluripotent stem cell (iPS)-derived brain endothelial cells (iPS-EC) were exposed to BME, VME and control sera and CSF and transendothelial electrical resistance (TEER in Ω) and capacitance (in nanofarad, nF) were measured as readout parameters of barrier function and endothelial monolayer integrity, respectively. **Figure 5** shows that both TEER (measured at 1000 Hz) and capacitance (measured at 64,000 Hz) were significantly differentially regulated in a time-dependent manner when exposed to pathogen-specific serum or CSF. TEER values reflect the transcellular barrier properties of the endothelial monolayer (**Fig 5A+B**), with the lower the TEER values, the more the barrier function is disturbed and vice versa. Capacitance reflects the intact integrity of the endothelial monolayer (**Fig 5C+D**), with the higher the capacitance values, the more patchy the monolayer and vice versa. CSF produced delayed effects with robust down-regulation of TEER in VME and BME at 4 and 24 h exposure compared to controls (**Fig 5B**), while capacitance is up-regulated late at 24 h in both VME and BME compared to control (**Fig 5D**). Serum samples produced similar but temporally different effects, but with significant early and intermediate downregulation of TEER already at 30 min and still at 4 h, but only in BME (**Fig 5A**). These effects disappeared after 24 h, TEER recovered and no differences were observed in either VME or BME compared to healthy controls. Capacitance was upregulated by serum samples after 4 h and 24 h and by CSF after 24 h (**Fig 5C+D**). It should be noted that CSF exposure generally increases barrier tightness, whereas serum exposure has the opposite effect (compare control values at different time points).

**Fig 5.**
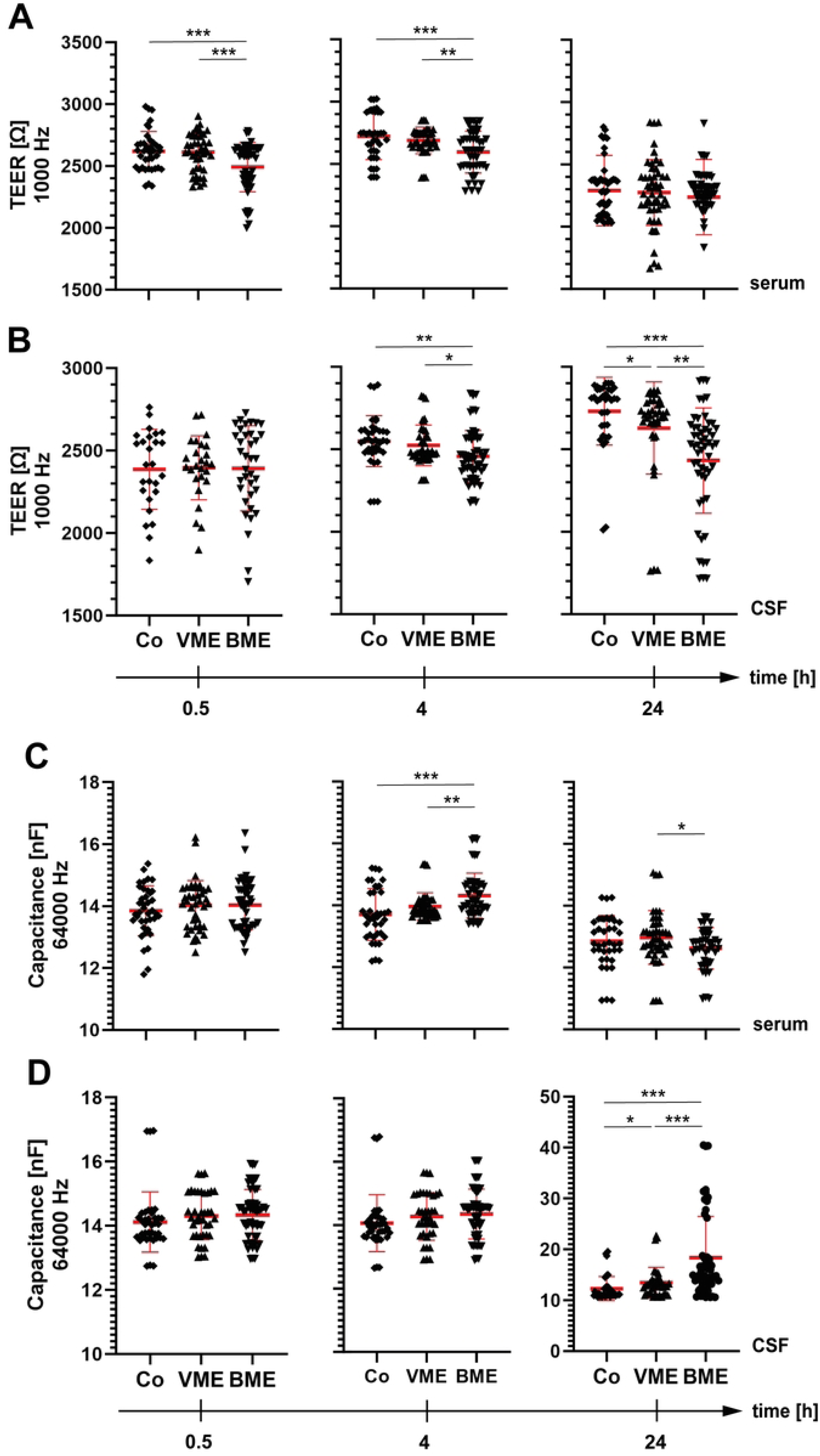
Regulation of BBB properties in response to pathogens. iPSC-induced brain endothelial cells (iP-EC) were grown in a monolayer on 96-well gold electrode array plates. iP-EC were exposed to serum (A, C) or cerebrospinal fluid (B, D) and TEER (A+B) and capacitance (C+D) were measured using the ECIS array system and are shown for three time points (30 min, 4 h, 24 h). TEER values are a marker of tight junction function, while capacitance, measured simultaneously, is a marker of overall intact endothelial monolayer integrity. Note that the higher the capacitance, the more leaky the monolayer, and the lower the TEER, the less tight the monolayer. A + B, show the pathogen-dependent effects of VME and BME, respectively, on tightness compared to healthy controls. C + D, show simultaneous regulation of capacitance. Legend: TEER, transendothelial electrical resistance measured at 1000 Hz and expressed in ohms (Ω • cm^2^); capacitance measured at 64 000 Hz and expressed in nanofarad (nF); VME, viral meningoencephalitis; BME, bacterial meningoencephalitis.

By dichotomising the readout parameters into favourable (RS ≤ 4) and unfavourable (RS > 4) ME patients, as described for Fig. 3 and 4, significant differences in BBB regulation could be demonstrated in both CSF- and serum-treated iPS-EC (**Fig 6**). In both serum- and CSF-treated cells, there was a significant downregulation of TEER after short-term exposure (30 min) in the unfavourable subgroup (**Fig 6A+B**), which was still present after 4 h of serum exposure (**Fig 6A**). In both CSF and serum, the effects were no longer present after 24 h. Capacitance was not significantly regulated by any treatment at any time (**Fig. 6C+D**).

**Fig 6.**
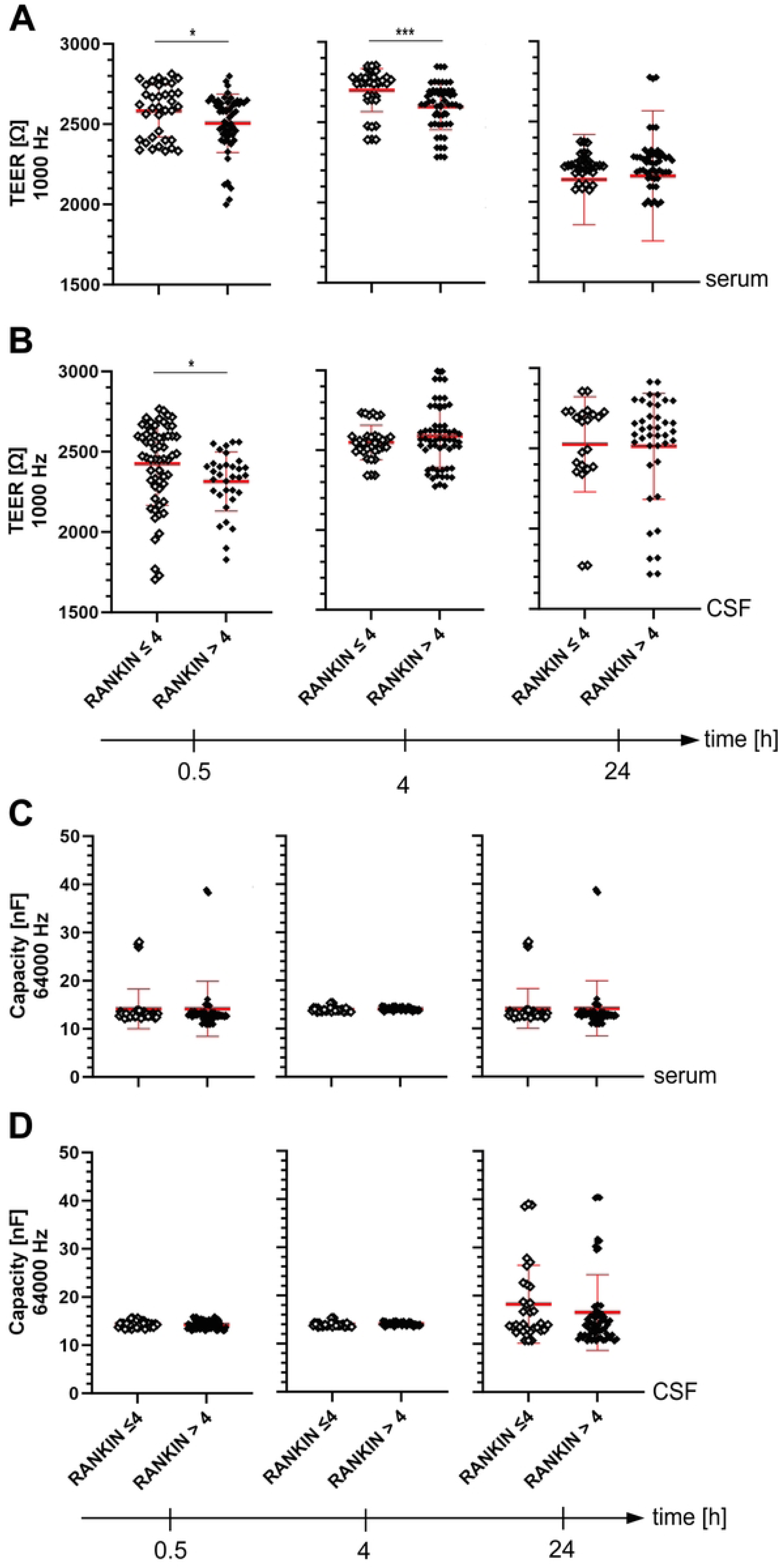
Regulation of BBB properties as a function of host (patient) disability. Shown is the regulation of TEER (A+B) and capacitance (C+D) of iP-EC cultures exposed to serum (A, C) or CSF (B, D) from patients with favourable (mRS ≤ 4) or unfavourable (mRS > 4) course of infection. RANKIN, modified RANKIN scale (mRS) for the measurement of disability (ordinal scale from 0 to 6, where 0 = not disabled at all, 6 = dead and further subdivided into 1-2 mild, 3-4 moderate and 5-6 severe). For general experimental design and further legends, see also Figure 5.

### Brain endothelial caspase activity

Caspase activity was analysed as a marker to quantify apoptosis in endothelial cells. Therefore, iP-EC were exposed to patient CSF or serum and caspase 3/7 activity was measured in cell lysates but also directly in patient’s serum and CSF samples. **Figure 7** shows caspase 3/7 activity measured directly in patient’s CSF samples (**Fig 7A-C**) and in exposed iP-EC lysates (**Fig 7D-G**). Caspase activity was significantly upregulated in iP-EC exposed to CSF from both VME and BME compared to healthy controls (**Fig 7E**). In contrast, serum from both BME and VME caused a significant downregulation of caspase activity in endothelial cells (**Fig 7D**). Looking at the effects on iPSCs in the dichotomised groups, CSF samples from patients with an unfavourable disease course (mRS > 4) induced more caspase activity in iP-EC than those from patients with less severe infections (**Fig 7G**); again, serum had no significant effect (**Fig 7F**).

**Fig 7.**
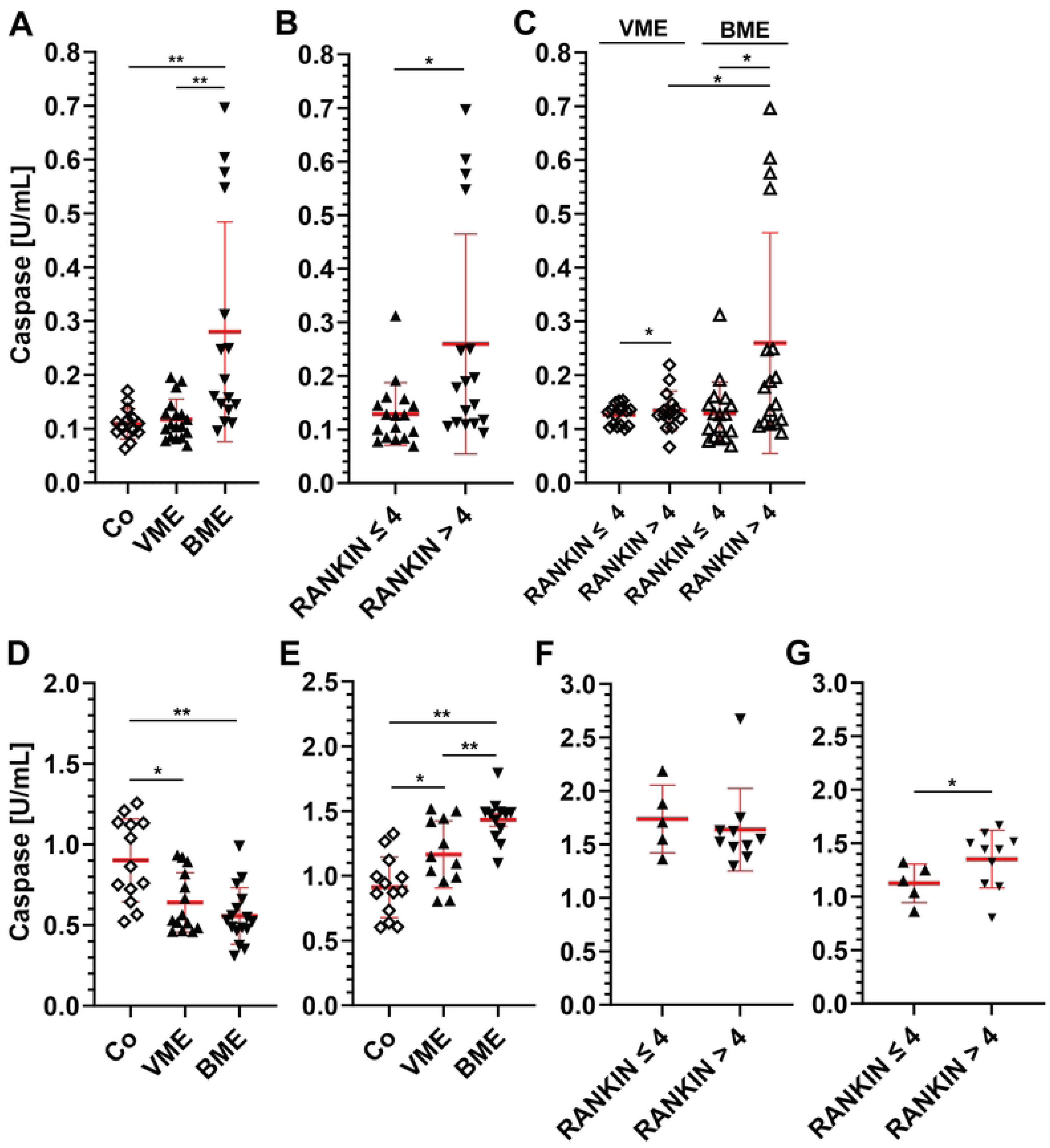
Endothelial cell death marker in meningoencephalitis. Caspase 3/7 activity was measured by cleavage of a luminogenic substrate Z-DEVD aminoluciferin directly in patient’s CSF samples (A-C) or in iP-EC lysates from cultures that were exposed to patient’s serum (D, F) or CSF samples for 24 h (E, G). Legend: iP-EC, iPSC-induced brain endothelial cells; VME, viral meningoencephalitis; BME, bacterial meningoencephalitis; RANKIN, modified RANKIN scale (mRS) for the measurement of disability (ordinal scale from 0 to 6, where 0 = not disabled at all, 6 = dead and further subdivided into 1-2 mild, 3-4 moderate and 5-6 severe).

Intrinsic caspase 3/7 activity could be also measured directly in the CSF and serum samples, but at much lower levels (factor 6 lower; compare **Fig 7A-C** and **Fig 7D-G**). A pathogen-specific caspase activity could only be detected in the CSF samples (**Fig 7A**), whereas this pattern was not observed in the serum samples (not shown). However, a significant increase in caspase activity was also detected in CSF samples from patients with an unfavourable course of infection (mRS > 4) (**Fig 7B**), which was also observed when the VME and BME subgroups were considered separately (**Fig 7C**). CSF from VME and BME patients caused a significant induction of caspase 3/7 activity in iP-EC (**Fig 7E**) as well as CSF from patients with unfavourable outcome increases caspase 3/7 activity compared to those with favourable outcome (**Fig 7G**).

### Regulation of endothelial drug transporter function

Having shown that CSF and serum from ME can specifically regulate blood-brain barrier function *in vitro* and induce endothelium derived caspase activity, we next investigated brain-specific regulation of endothelial drug transporters. For this purpose, patient samples were exposed to either hCMEC cells, which strongly express p-glycoprotein (P-gp), or MDKII-BCRP cells, which express breast cancer resistance protein. Both P-gp and BCRP belong to the ATP-binding cassette transporter superfamily and are expressed in human brain endothelium [30, 31]. **Figure 8** shows the regulation of P-gp and **Figure 9** the regulation of BCRP transporter function in response to CSF and serum after short-term (2 h) and long-term (24 h) exposure. Short-term responses (2 h) should reflect non-gene regulatory direct responses of transporter activity, whereas long-term responses (24 h) may also include gene regulatory effects. However, we did not pursue this further.

**Fig 8.**
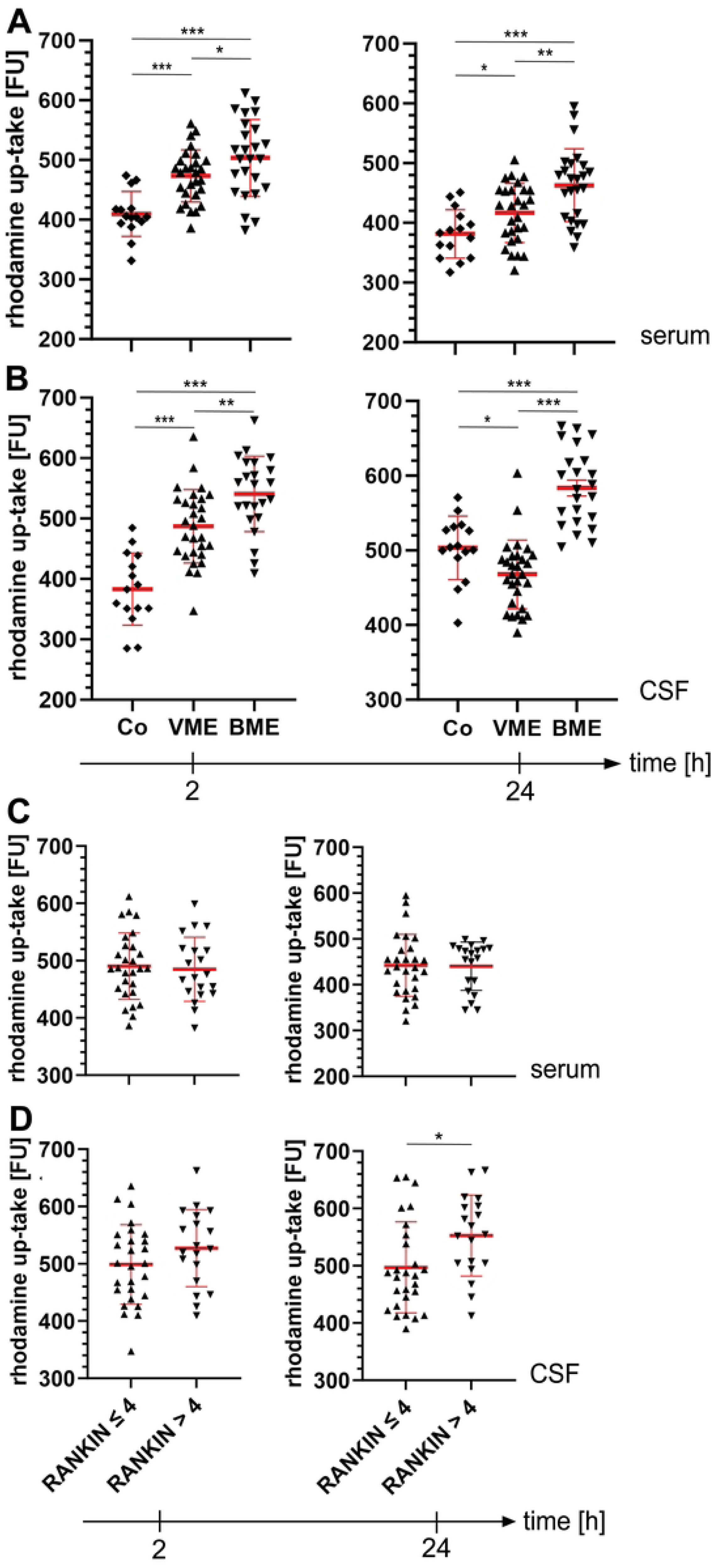
P-gp drug transporter activity in response to pathogens and as a function of host (patient) disability. hCMEC/D3 cell-line constitutively expressing P-gp were grown to confluence for 4 days in 96-well plates and exposed to either patient’s serum or CSF at the indicated time points. Intracellular accumulation of the specific P-gp substrate Rhodamine 123, as a measure of transporter inhibition, was measured in cell lysates at specific wavelengths (λ absorbance at 485 nm, λ emission at 535 nm). (A) shows the uptake after exposure to patient serum and (B) to CSF in VME and BME compared to healthy controls. (C, D) shows the respective rhodamine uptake according to the severity of infection, dichotomised into favourable (mRS ≤ 4) and unfavourable (mRS > 4) subgroups. Legend: FU, fluorescence unit; P-gp, P-glycoprotein; VME, viral meningoencephalitis; BME, bacterial meningoencephalitis; RANKIN, modified RANKIN scale (mRS) for the measurement of disability (ordinal scale from 0 to 6, where 0 = not disabled at all, 6 = dead and further subdivided into 1-2 mild, 3-4 moderate and 5-6 severe).

**Fig 9.**
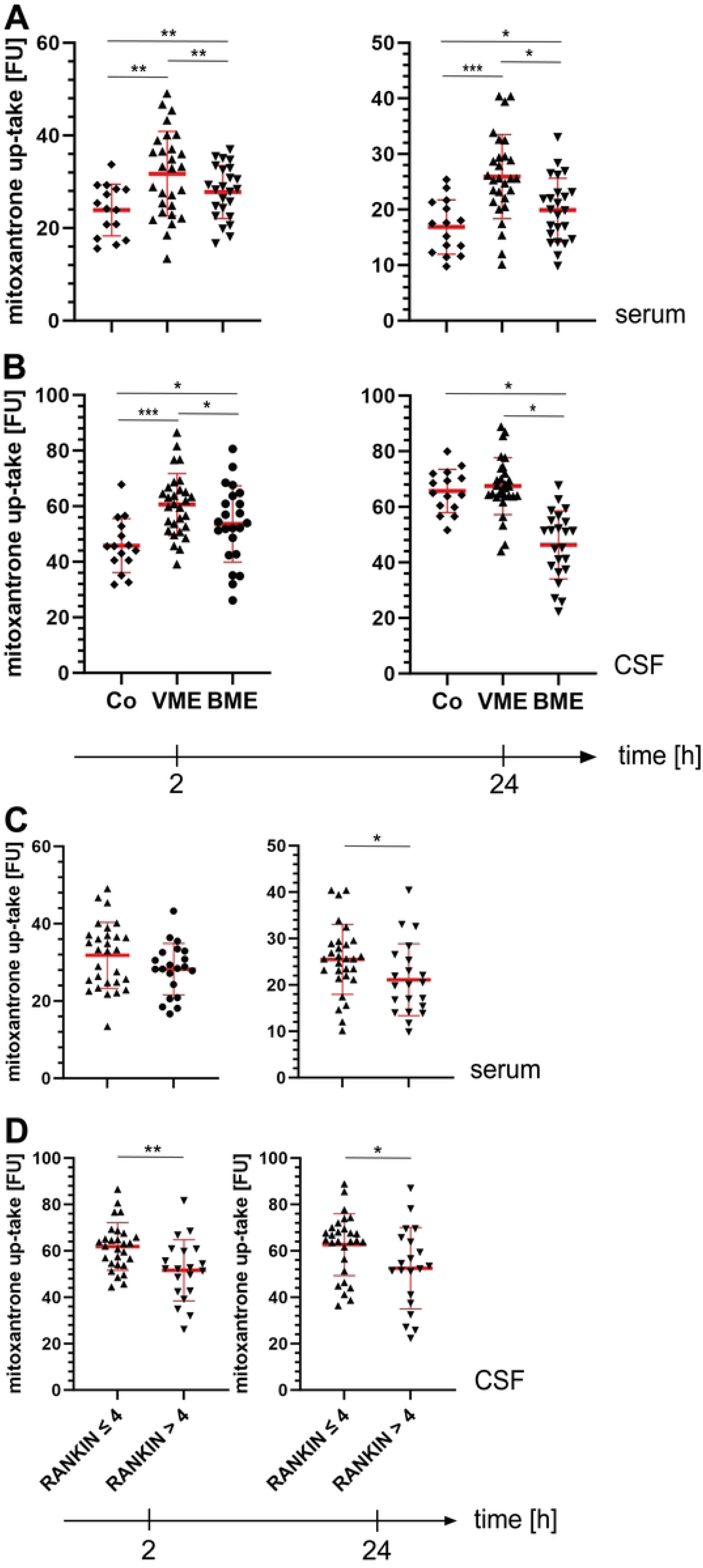
BCRP drug transporter activity in response to pathogens and as a function of host (patient) disability. MDKII-BCRP cell-line constitutively expressing BCRP were grown to confluence for 4 days in 96-well plates and exposed to either patient’s serum or CSF at the indicated time points. Intracellular accumulation of the BCRP substrate Mitoxantrone, as a measure of transporter inhibition, was measured in cell lysates at specific wavelengths (λ absorbance at 610 nm, λ emission at 685 nm). (A) shows the uptake after exposure to patient serum and (B) to CSF in VME and BME compared to healthy controls. (C, D) shows the respective mitoxantrone uptake according to the severity of infection, dichotomised into favourable (mRS ≤ 4) and unfavourable (mRS > 4) subgroups. For further legend details, see Fig 8.

P-gp transporter activity, measured by the accumulation of Rhodamine as its specific substrate, showed a significant accumulation of the specific fluorescent substrate Rhodamine in hCMEC cells after stimulation with both CSF and serum from VME and BME patients, even after short-term and long-term exposure, indicating P-gp inhibition. However, in the VME group, this increase disappeared after 24 h exposure of the cells to CSF, indicating rather P-gp activation. (**Fig 8A+B**).

The results for BCRP activity, assessed by the accumulation of the specific substrate Mitoxantrone in MDKII BCRP cells, were in principle similar to those for P-gp, but differed in some aspects (**Fig 9**). The inhibitory effects of CSF and serum were stronger after exposure to VME samples compared to those exposed to BME samples (**Fig 9 A+B**). Furthermore, BME caused activation of BCRP in CSF-exposed cells after long-term exposure, as evidenced by reduced mitoxatrone accumulation after 24 hours (**Fig 9B**).

Looking again at the dichotomised groups, an inhibition of the transporter activity for P-gp in the severely affected group (mRS > 4) was only seen in the CSF-exposed cells after long-term exposure; early CSF effects and serum effects in general could not be detected (**Fig 8C+D**). However, the effects on BCRP activity were different. Both serum and CSF caused a decrease in Mitoxantrone concentration in the cells, which corresponded to BCRP activation in the severely affected group (mRS > 4) (**Fig 9C+D**).

## Discussion

Our profiling identified 36 (67%) of the 54 factors studied that were significantly up- or down-regulated compared to healthy controls. Of these, CCL2, CCL3, CCL27, IL-3, IL-13, IL-16, TNF-β, and NGF were identified as previously undescribed candidates in BME, and IL-7, IL-9, FST, FGF-2, HGF, CLEC11A and VCAM-1 were identified as such in VME (**Table 2**). The remaining differentially regulated factors we identified have already been published [32–41]. Thus, our ability to reproduce a subset of known differentially regulated factors serves as an internal positive control for our overall study in terms of quality and validation of our novel findings. Furthermore, GSEA identifies downstream signaling pathways of enriched viral and bacterial-associated factors that have already been identified, such as fever induction or LPS-mediated signaling in BME or positive regulation of MHC class II proteins in VME (not shown). Many of the identified and differentially regulated factors correlate significantly, either positively or negatively, with the severity of the infection as measured by the mRS as a measure of patient disability (**S1 Fig.**). In addition, QAlb, as a measure of blood-CSF barrier dysfunction, positively correlates with disability in our cohort. QAlb has previously been described as a negative prognostic factor for the outcome of pathogen-associated meningoencephalitis [42, 43]. Furthermore, CSF glucose levels are inversely correlated with mRS score in BME. CSF glucose levels as negative prognostic factors in BME have already been described [43, 44]. Thus, we can replicate known findings at the level of barrier dysfunction in both bacterial and viral meningoencephalitis.

We were particularly interested in understanding whether inflammation in the brain responds to bacterial or viral infection not only with upregulation, but also with down-regulation compared to the healthy state. This approach offers the possibility of identifying potential factors targeted by pathogen virulence factors that could act as host protective factors to maintain neuronal homeostasis under healthy conditions (**Fig 1 + 2**). Thus, in BME, predominantly growth factors that can exert neuroprotection and -regeneration were identified to be down-regulated (**Fig 1A**) [45–53]. In addition, FGF-2 and NGF work synergistically in exerting their protective effects [46, 54]. GSEA (**Fig 1B**: cluster 2) revealed neurotrophin- and FGF-2 signaling and their downstream MAP kinase and phosphatidylinositol signaling pathway - directly affecting neuronal cell growth, differentiation, development, and survival - as putative attenuated pathways in BME. In addition, FGF-2, PDGFB, and VEGFA enhance neurotrophic pathways via BDNF, one of the strongest neurotrophic factors and mediator of neuroplasticity [55, 56]. Accordingly, these neurotrophic factors are strong inhibitor of pro-aptototic signals (**Fig 1B**: cluster 1). Interestingly, all factors also convert into Notch1-signaling (**Fig 1B**: cluster 3), which is also involved in neurogenesis and neuroprotection [57, 58]. Furthermore, Notch-1 signaling is involved in controlling reactive astrocytes, which are an important source for neurotrophic support [59]. Moreover, reactive astrocytes have a pivotal function in antigen presentation and regulation of pro- and anti-inflammatory effects in brain inflammation [60]. Similarly, GSEA revealed that the same negatively regulated factors identified here can activate a signaling cascade via IL10 that converts into SOCS3 signaling, known as a negative regulator of inflammation (**Fig 1B**: cluster 4). In addition, SOCS3 has been described to inhibit neuronal cell death independently of classical neurotrophic signaling [61, 62]. Last but not least, expression of matrix-metalloproteases (MMP) seem to be affected in BME (**Fig 1B**, cluster 5). MMPs have an important function in processing and consecutively activating growth factors such as FGF-2 [63]. In summary, the loss of trophic support at multiple levels during bacterial infection of the brain may lead to critical changes in homeostasis and may contribute significantly to neuronal loss.

In VME, a different group of interleukins and growth factors are downregulated, with the exception of VEGFA and HGF, which were found to be downregulated in both VME and BME compared to healthy conditions (**Fig 2A**). GSEA revealed that similar but also different downstream signaling pathways are targeted, for example the positive regulation of BDNF (**Fig 2B**: cluster 1-3). However, GSEA suggests that CD40L/STAT signaling may be less active in VME compared to healthy controls. Both play important roles in the viral adaptive and innate immune response via antigen presentation, cell-mediated Th1 (CD40L) and toll-like receptor-mediated responses [64, 65]. Thus, in contrast to peripheral tissues, the brain may be slower to respond to these initial antiviral steps. In addition, the downregulated factors appear to target antiviral pathways via CD274/CD44 and chemokines (**Fig 2B**: cluster 4+5). CD272 (syn. B- and T-lymphocyte attenuator or BTLA) has been implicated in hepatitis B and herpes virus pathology [66]. CD272 deficiency can impair effector CD8 T cells in a vaccinia virus infection model [67]. Together with herpes virus entry mediator (HVEM), BTLA appears to play a critical role in immune tolerance [66]. CD44 can reciprocally induce CXCL10 via interaction with TLR on virus-infected cells and support virus-specific cytotoxic T lymphocyte migration [68]. Consequently, we can show that CXC motif chemokine receptor signaling is less active in VME (**Fig 2B**: cluster 5). Less is known about CXCL signaling in human VME. However, CXCLR4 antagonism appears to improve survival in West Nile virus encephalitis [69]. Interestingly, CXCL3-deficient mice showed improved viral clearance and reduced neuroinflammation in experimental herpes encephalitis [70]. Notably, 12 of the 23 viruses in our VME subgroup were members of the herpes virus family. Thus, CXCL signaling may play a critical role in the parenchymal infiltration of T lymphocytes and subsequent viral clearance. Last but not least, tight junction protein (TJP) activity might be dysregulated in VME (**Fig 2B**, cluster 6). TJP play a role in cellular virus spread and tropism [71, 72]. Therefore, temporary down-regulation of tight junction proteins on the brain endothelium may be beneficial to prevent virus transmigration across the blood-brain barrier. In conclusion, in VME the loss of trophic signals may also promote an environment that leads to neuronal loss. In addition, antiviral and immune tolerance mechanisms may be incompletely activated in the brain.

We were also interested in identifying host (patients) factors that may contribute to the severity of infection and level of disability. By dichotomising the level of disability regardless of whether patients had VME or BME, we were able to identify specific factors that are differentially regulated with either a favourable (mRS ≤ 4) or unfavourable (mRS > 4) course of infection (**Table 3**). All identified factors correlated, either positively or negatively, not only with the dichotomisation but also with the grade of mRS (**Fig 3**). However, it was not possible to perform GSEA because both subgroups and total number included, respectively, were too small. Nevertheless, based on an extended structured Boolean literature search, putative downstream pathways could be described for both the favourable (mRS ≤ 4) and unfavourable (mRS > 4) subgroups and are summarised in **Fig 4** (for detailed Boolean literature search, see also **Methods Appendix S1** and **S3 Table**). Patients belonging to the favourable (mRS ≤ 4) subgroup appear to preferentially activate downstream signaling pathways of neuroprotection and apoptosis and may better regulate the termination of the immune response (**Fig 4A**). For example, IL-10 and SOCS3 have been shown to be critical in terminating the inflammatory response in *St. pneumonia* and *Mycobacterium tuberculosis* infections; note that *St. pneumonia* contributes significantly to our BME group [73, 74]. The role of CD274/PD-L1 in anti-inflammatory signaling appears to be pathogen-dependent, as data suggest a more negative role, except in HSV-1 infection where it appears to contribute to viral resistance; in particular, members of the herpes virus family are a major viral representative in our study [75]. In addition, CSF3, HGF, and SCF1 can exert neuroprotective effects and can induce BDNF, CNTF, and NGF [76, 77]. All converge on PI3K/AKT and MAPK/ERK pathways to regulate neuronal survival and plasticity [55].

Finally, IL-6, HGF, CSF3 and SCF have been implicated in the regulation of apoptosis via p53, Bcl-2, Bcl-xL, Bax and caspase 3 in neuronal lesion models [78, 79]. The data derived for apoptosis must be interpreted with caution, as both anti-apoptotic and pro-apoptotic pathways can be targeted (**Fig 4A**). In addition, many of the Boolean literature search results are from oncology, so an uncritical translation to brain infections is not readily possible. However, it has recently been shown that p53 may be critically involved in the clearance of bacterial pathogens [80, 81]. Thus, both pathways may promote a favourable outcome in brain infection, as the termination of an immune response depends on the apoptotic clearance of immune cells, and activated anti-apoptotic pathways are essential to mediate neuroprotection.

However, in the unfavourable subgroup (mRS > 4), IFNγ-regulated genes, CTL/NK cell activation and neurotoxicity pathways appear to be more strongly activated (**Fig 4B**). As critical signaling molecules in infection, JAK-STAT-dependent signaling plays a key role in pathogen defence [82]. A certain level of STAT activity may be critical for an appropriate immune response to a given pathogen [83]. Interestingly, two factors, IL-3 and IL-12, which were found to be upregulated in the unfavourable subgroup, can negatively regulate SOCS1, an important negative feedback suppressor of the JAK1/STAT1 IFN pathway, potentially leading to an unleashed immune response [84–86]. In addition, the observed increased IL-12 and IL-15 activity may be indicative of increased CTL and NKC activity [87–90]. Together with the observed elevated levels of CXCL9 and CXCL12, both of which can exert neurotoxic effects [91–95], the pathways identified here may create a cytotoxic environment in the brain of patients with unfavourable infection course.

The integrity of the blood-brain barrier (BBB) is central to intact brain function in terms of oxygen and nutrient delivery, as well as of metabolite, drug and toxin transport. Disruption of the BBB has been associated with several inflammatory diseases such as septic encephalopathy, cerebral malaria or, more recently, Covid-19, and can increase morbidity and mortality [96–98]. The expression of members of the ABC drug transporter family at the apical membrane of the brain endothelium (and in cellular organelles such as mitochondria) is an important element of BBB function, forming a selective and active transport barrier. Among these, P-gp and BCRP have been identified to limit the penetration of drugs and toxins into the brain. We investigated the effects of CSF and serum from ME patients on brain endothelial function in vitro using a human iPSC-derived BBB model and cell lines selectively expressing the drug transporters P-gp and BCRP. We could demonstrate significant effects at several cellular and molecular levels of the BBB, such as regulation of endothelial tightness measured as TEER, endothelial layer integrity measured as capacitance, disruption of transporter function measured as P-gp and BCRP activity, and at the level of inducible endothelial cell death measured as caspase 3/7 activity (**Fig 5-9**). These effects were demonstrated for both groups of pathogens examined (viruses and bacteria) and depending on the severity of the infection, which suggests host-(patients-) dependent effects.

At the level of BBB tightness, we observed differential effects when serum and CSF were considered separately. We also observed differential effects depending on the temporal dynamics. Both serum and CSF showed rapid effects with down-regulation of TEER within 30 min up to 4 h, most likely direct receptor-mediated effects at the level of tight junction proteins. The effects of CSF were generally stronger than those of serum and were still present at 24 h (**Fig 5A+B**). Proinflammatory cytokines such as IFN-γ, IL-13, IL-16 or IL-17, which we found to be elevated in either BME or VME patients, are known to rapidly downregulate tightness by phosphorylating tight junction proteins via their receptor-mediated signaling pathways [99–101]. However, after 24 h only the CSF effects persisted, whereas the serum effects disappeared (**Fig 5A+B**). Unlike TEER, which changes rapidly upon stimulation, cell layer capacitance reflects the overall functional and morphological integrity of the endothelial cell. Our data showed that after prolonged exposure to CSF, endothelial cell layer capacitance increased in both the viral and bacterial groups, indicating increasing disruption of the endothelial monolayer beyond the rapid molecular effects on TEER regulation (**Fig 5D**). The loss of cells within the endothelial monolayer is further supported by our observation of a concomitant increase in caspase 3/7 activity in endothelial cells exposed to BME and VME CSF (**Fig 7E**). Accordingly, IL-1 and IFN-γ, both of which have been shown to be upregulated in either BME or VME (**Table 2**), can induce endothelial apoptosis. In addition, endothelial apoptosis is also directly mediated by members of the TNF receptor superfamily CD40/CD40L (CD154), which are upregulated by IL-1 and IFN-γ [102–105].

Looking again at the cellular effects on the BBB depending on the severity of the infection, our results show that the BBB seems to be more dysregulated after exposure to CSF and serum of severely affected patients (mRS > 4) (**Fig 6A+B**). These effects are further supported by our caspase 3/7 data. CSF from the more severely affected group leads to more pronounced caspase activity in endothelial cells (**Fig 7G**). Furthermore, we were able to confirm this observation by measuring caspase activity directly in the CSF samples. Again, we found higher caspase activity in the CSF samples of the more severely affected patients (**Fig 7B**), which can be traced back even further when the effects of VME and BME are considered separately (**Fig 7C**). In COVID-19, deterioration of the BBB has been intensively discussed as a putative mechanism for neurological symptoms [98]. Recently, elevated MMP levels in COVID-19 have been shown to be a marker of BBB dysfunction. These findings are consistent with our analysis of signaling pathways for both viral and bacterial ME, which identified both MMPs and tight junction proteins (latter in VME only) as downstream targets (**Fig 1B** and **Fig 2B**). In addition, a blood-brain barrier disruption and specific inflammatory profile that partially overlaps with the inflammatory profile in our ME cohorts has recently been demonstrated in Post-Covid patients. [106].

Looking at a more functional aspect of the brain endothelium, we were also able to show short- and long-term effects on drug transporters upon stimulation with patients CSF or serum. P-gp and BCRP activity was inhibited shortly after 2 hours, regardless of whether the cells were exposed to serum or CSF from VME or BME patients (**Fig 8** and **Fig 9**). These effects are generally maintained after long-term exposure. However, there are two differences. CSF from VME patients induces P-gp activity and vice versa, CSF from BME patients induces BCRP activity after 24 h. Little is known about the role of drug transporter function in the context of inflammation. However, there is evidence that P-gp is required to induce a full IFN type I response following infection with *Listeria monocytogenes* [107]. HIV pg120 can directly inhibit P-gp expression in astrocytes [108]. In addition, P-gp function has been shown to be essential for natural killer cell and T lymphocyte-mediated cytolysis [109]. Our data may support an immunoregulatory role for P-gp in addition to its classical transporter function. In general, however, the inflammation-induced inhibition of drug transporter activity can have important wider consequences. Inhibition can lead to accumulation of transporter substrates - such as drugs, but also endogenous and xenogenous toxins - in the brain, causing secondary brain damage. Accordingly, an increase in P-gp and BCRP substrates has been demonstrated in human inflammation [110]. Our observation that P-gp inhibition is more pronounced in the group of more severely affected ME patients may therefore be indicative (**Fig 8C+D**). However, the effect on BCRP was the opposite and the significance of this observation remains unclear (**Fig 9C+D**).

An intact neurovascular unit (NVU) is essential for maintaining homeostasis in the brain and has a direct impact on neuronal function and network stability. Although we did not study the NVU directly, many of the findings of the study can be extrapolated to possible effects at the level of the NVU. It has long been known that circulating inflammatory mediators during systemic inflammation, such as sepsis, induce multiple alterations in the BBB, ultimately leading clinically to delirium or septic encephalopathy [111]. Cerebral malaria is a well-studied prototypical systemic infection that has dramatic effects on brain function through inflammation of the BBB without actually infecting the brain itself [112]. Understanding of the effects on the BBB during direct infection of the brain and contributions to the pathophysiology of pathogen-mediated parenchymal damage on the one hand and inflammation-induced widespread effects on the other, and the ultimate consequences for patient morbidity and mortality, is still poorly understood. In bacterial meningitis, the mechanisms of bacterial invasion of the brain have been mainly studied using BBB models. Bacterial pathogens adhere to brain microvascular endothelial cells, which form the major component of the BBB, and can then cross via receptor-mediated transcytosis into the subendothelial parenchyma. This activates the NF-kB and Type 1 IFN signalling pathways in endothelial cells, leading to leukocyte transmigration across the endothelium. These initial steps are known as the “pathogenic triad” [113]. In viruses, neurotransmission depends on virus species-specific routes. For members of the alpha-herpes family, HSV-1 and VZV, transmission to the brain is still unclear and may depend on anterograde axonal transport from the site of latent infection (sensory ganglia) or via haematogenous spread, or both. [114]. However, for TBEV, several routes of infection have been postulated in addition to receptor-specific infection of the cerebral endothelium [115]. Note that HSV-1, VZV and TBEV are the main representatives of our VME subgroup. Systemic inflammation has significant consequences for the function of the brain at the level of the NVU as shown by changes in signaling of the cerebral endothelium, enhanced cellular traffic, increased solute permeability and direct damage to the endothelium [113]. However, it remains unclear how the infection affects the brain beyond its infection core and what the overall consequences of the NVU disruption might be. Our data show that in infectious meningoencephalitis there is multiple regulation of inflammatory mediators, but also of neurotrophic (growth) factors and neuroprotective interleukins, which may have effects beyond the endothelium itself. Using a Boolean search strategy, we attempted to relate the findings from our BBB model and the findings on regulated mediators to possible pathophysiological effects at the level of individual NVU components (**Fig 10, S4 Table**).

**Fig 10.**
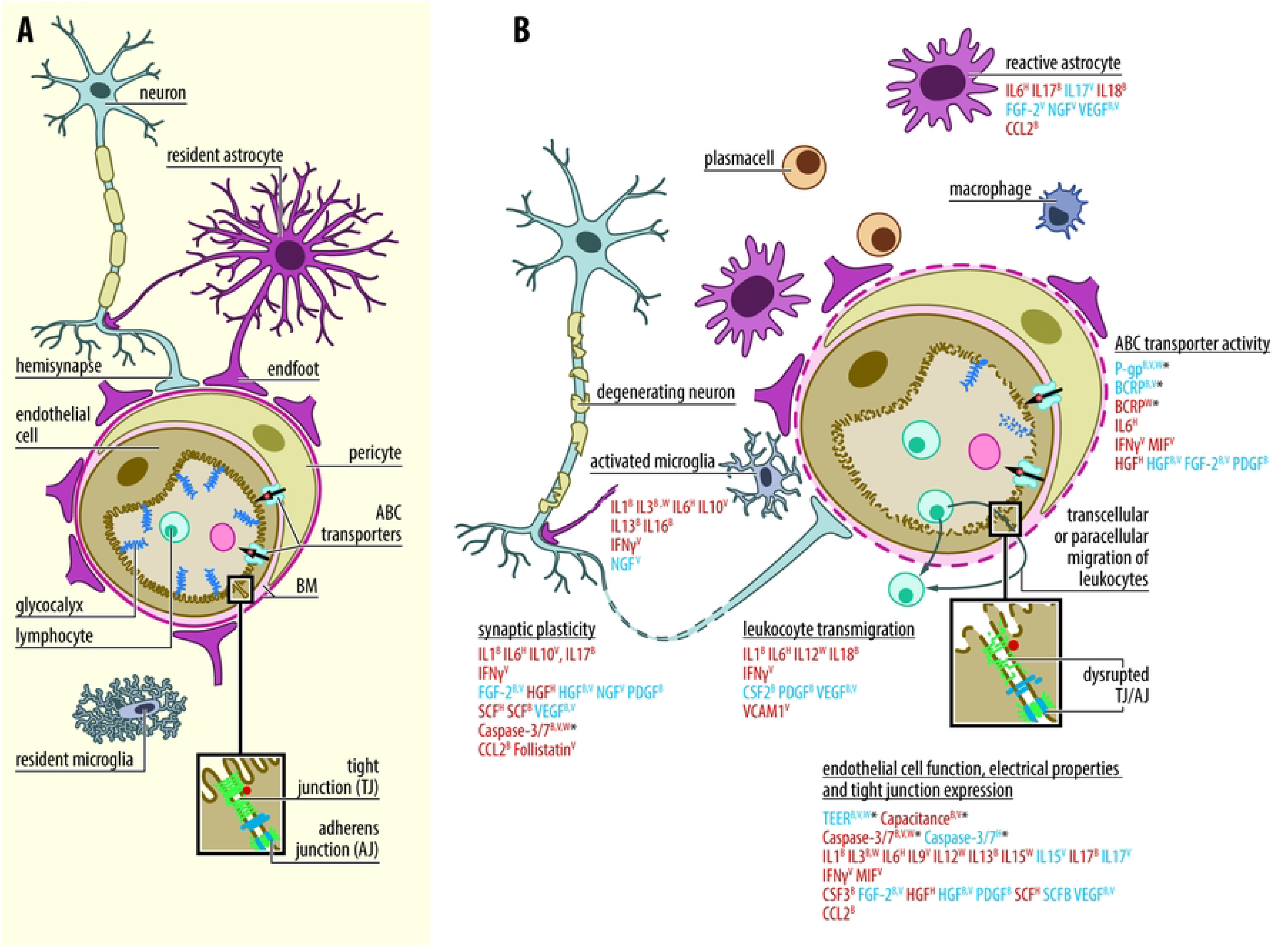
Impact of study findings at the level of the neurovascular unit. Schematic representation of the components of the neurovascular unit (NVU) under physiological conditions (A) and putative effects at different ultrastructural levels based on study findings in pathogen-associated meningoencephalitis (B). Legend: BM, basal membrane; factors/function in bold red, upregulated according to study results; factors/function in bold blue, downregulated according to study results; superscript B, regulated in bacterial infection; superscript V, regulated in viral infection; superscript W, regulated in patients with poor outcome; superscript H, regulated in patients with favourable outcome; *, direct effects measured in the IP-EC BBB model according to study. Note that the cell proportions are not to scale and the astrocyte endfeet are not shown fully wrapped around the BM for better visualisation. See S4 table for references.

Thus, downregulation of neurotrophic growth factors such as FGF-2, HGF, NGF, PDGF, SCF or VEGF, either in patients with poor outcome or in bacterial or viral meningoencephalitis, may have significant consequences for the function of the NVA at different ultrastructural levels. All these factors have more or less neuroprotective effects and promote synaptic plasticity but also stabilize electrical properties and tight junction function in brain endothelium [46, 116–125]. In addition, downregulation of these factors may alter drug transporter function and may (in part) reflect mediators of impaired BBB function, as we were able to demonstrate in our IP-EC model at the level of electrical properties and transporter function [126, 127]. Interestingly, all of the factors that we found to be upregulated in the subgroup of patients with a favourable outcome may also orchestrate the physiological stabilisation of NVU, whereas those, including chemokines and interleukins, that were found to be preferentially upregulated in the subgroup with a poor outcome may have the opposite effect and exert immune-related neurotoxicity (**Fig 4** and **Fig 10**). For example, simply overexpression of IL3 in transgenic mice leads to strong microglial and astrocyte activation and consequently to motor neuron disease [128]. IL12 is one of the key mediators of cerebral malaria, a prototypical BBB disorder, and mediates neurotoxic capacity via IL12-p40-iNOS signaling [129, 130]. Chemokines such as CXCL9 or CXC12 released by microglia or macrophages can lead to impaired synaptic plasticity and to learning and memory deficits [131]. Interestingly, it has recently been shown that a disruption in P-gp function, which is downregulated particularly in our BME group, but also in the subgroup with severe infection, leads to accumulation of aldosterone in the brain and anxiety-like behaviour [132, 133]. Recent data identify similar immune-mediated BBB dysfunction in Covid-19-associated neurological symptoms and in post-Covid syndrome patients with increased immune-mediated neurotoxicity [106, 134, 135]. As a result, all of these adverse conditions seen in our patients may alter NVU function and contribute to increased morbidity and ultimately increased mortality.

Finally, the limitations of our study need to be discussed. These include the limited number of patients allocated to the experimental groups and the healthy controls. However, despite the limited number of cases, we were able to find strong and significant effects between the different pathogen groups (BME versus VME) and within the subgroup analysis (favourable versus unfavourable). Secondly, we only included a limited number of different pathogen species. Therefore, the number of different pathogen species needs to be increased to cover a wider range of host-specific immune responses. Thirdly, due to our study design, we were not able to attribute a differentially regulated factor and its derived putative signaling pathway to either an immune or neuronal cell origin. This could be better investigated by studying both the soluble CSF inflammome and the transcriptome of simultaneously isolated CSF leukocytes. Finally, the putative pathways and associated biological responses as a function of infection severity shown in **Figure 4** should be interpreted with caution, as they are based only on a structured Boolean literature search and not on an algorithm as used by GSEA. In this way, GSEA or comparable algorithms would be able to weight the effect of individual factors within a given network. Therefore, we would like to emphasise that the pathways and the associated downstream biological output responses are scattered in nature. Further in-depth analysis of the inflammome in specific pathogen-induced meningoencephalitis is required to substantiate the relevance of the pathways found.

## Conclusion

We have identified novel and specific differentially regulated inflammatory pathways associated with bacterial and viral infections of the brain. On the host side, patients with a favourable course of infection may benefit from a more balanced immune response that can more effectively eradicate the causative pathogen and appropriately regulate and terminate inflammation in time to reduce neuronal damage caused by excessive inflammation. Similarly, increased neurotrophic support may protect against neuronal death and promote regeneration. An inverse environment, characterised by an unleashed immune response and increased cytotoxic activity, may be present in patients with severe course of infection. Complementary analyses reveal a disrupted blood-brain barrier function in vitro - both pathogen- and host-dependent - in terms of endothelial tightness, drug transporter activity and endothelial cell death. Further in-depth network analysis will help to narrow down the critical key signals in misdirected immune responses and should enable the crucial pathways to be elucidated in more detail. Once these patterns are elucidated, there is an opportunity to intervene with specific therapeutic strategies beyond established anti-infective treatment to contain unleashed immune responses or to boost inappropriately weak responses [136].

## Materials and Methods

### Study design, patient selection and ethics

We conducted an observational cohort study to investigate pathogen- and host-dependent inflammatory patterns and blood-brain barrier (BBB) regulation in pathogen-specific meningoencephalitis. Patients with viral (VME) and bacterial (BME) meningoencephalitis and controls were enrolled at the Department of Neurology, Nordwest Hospital, Frankfurt, or at the Department of Neurology, University of Heidelberg. Patients were either prospectively enrolled or retrospectively selected from the hospital information system. For prospectively enrolled patients, CSF and blood (serum) samples were obtained during a routine lumbar puncture for clinical diagnosis. For retrospectively selected patients, frozen CSF and serum were obtained from our CSF biobank. Control CSF and serum samples were obtained from healthy subjects, e.g. to exclude subarachnoid haemorrhage. The clinical data set included the modified RANKIN scale (mRS) [26]. Immunosuppressed patients were excluded. The study was conducted in accordance with the tenets of the Declaration of Helsinki, approved by the Ethics Committee of the Landesärztekammer Frankfurt, Hessen, and the University of Heidelberg, and informed consent was obtained.

### Quantification of cytokines, chemokines and growth factors

Frozen CSF/serum sample-pairs were thawed (ice) and processed to analyze inflammatory factors on the same day using the Bio-Plex Systems® (Bioplex Pro Assay^TM^®, Bio-Rad Laboratories GmbH, Munich, Germany) and were measured with the Luminex 200 system as reported previously [137]. The following multiplex bead arrays were investigated: Bio-Plex Pro™ Human Cytokine 27-Plex Panel (Bio-Plex Pro Human Cyto 27-Plex Panel, M500KCAF0Y, Bio-Rad Laboratories GmbH, Munich, Germany), Bio-Plex Pro™ Human Cytokine 21-Plex Panel (Bio-Plex Pro Human Cyto 21-Plex Panel, MF005KMII, Bio-Rad Laboratories GmbH, Munich, Germany), and Bio-Plex Pro™ Human Angiogenesis 9-Plex (Bio-Plex Pro Human Angiogenesis, 171A4S11M, Bio-Rad Laboratories GmbH, Munich, Germany) (for overview of investigated factors, see **S1 Table**). Dilutions of samples: serum samples were diluted in sample diluent (1:20) according to the manufacturer’s guidelines, whereas CSF dilution was empirically tested and was finally diluted 1:2 in sample diluent. The specific intrathecal proportion of a given factor was calculated by factoring in the CSF/serum albumin quotient as: 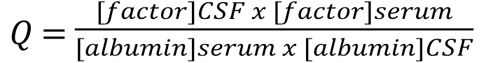 [138–140]. Raw results for each factor were originally reported in pg/ml. In all figures, factor concentrations are expressed as Q [factor/albumin] unless otherwise stated in the legend.

### Analysis of blood-brain barrier function in vitro

As a model, we studied human brain endothelial cells (iPS-EC) derived from pluripotent inducible stem cells (iPSC). SBAD03 iPSC were derived from human skin biopsy fibroblasts following signed informed consent, with approval from the UK NHS Research Ethics Committee (REC: 13/SC/0179) and were derived as part of the IMI-EU sponsored StemBANCC consortium. iPSC generation was performed using the CytoTune-iPS 2.0 Sendai Reprogramming Kit (A16517) from Thermo Fisher Scientific (Waltham, MA), as previously described [141, 142]. Briefly, 7.5x10^3^ iPSC/cm2 were seeded on Matrigel coated (Corning, Wiesbaden, Germany) 6 well plates and maintained in mTeR™1 medium (Stemcell Technologies, Grenoble, France) for three to four days (day -3-4) until having reached a density of 2.5-3.5x10^4^ cells/ cm2. Next differentiation of iPSC into neuronal and endothelial cells took place in the presence of unconditioned medium containing 100µM β-mercaptoethanol for 6 days. Expansion into BBB endothelial cells (hiPS-EC) was induced by cultivating cultures in endothelial serum free medium (ESM, Life Technologies, Darmstadt, Germany) supplemented with B27, FGF-2 (20ng/mL) and retinoic acid (10µM) for another two days. At day 8, purification of hiPS-EC was reached by subculturing the splitted cells on a selection-collagenIV/fibronectin-matrix (Thermo Fisher Scientific, Ulm, Germany; Merck, Darmstadt, Germany) already in culture plates for a given finale experiments. Experiments finally took place at day 10 of the differentiation and selection procedure. Patient’s sera and CSF were analysed in 96 well array gold grid electrode plates (96W20idf, Applied Biophysics, Ibidi, Gräfeling, Germany) at 100µL ECM/well with 25% (v/v) for serum and 50% (v/v) for CSF. Resistance of the hiPS-ECs were measured with the ECIS® ZØ instrument (Applied Biophysics, purchased from Ibidi, Gräfeling, Germany). Confluency of cultures starting with final seeding was monitored and validated using impedance and capacitance analyses. All experimental values are given within the figures as resistance (Ω • cm^2^) or capacitance (nF) in absolute values at steady state cultures.

### Drug transporter up-take and regulation

For the *in vitro* analysis of P-gp and BCRP drug transporter function, hCMEC/D3 and MDKII-BCRP cell-line, respectively, were investigated. Cell-lines were cultured and maintained as described elsewhere [143, 144]. For the experimental set-up, cell-lines were seeded into 96 flat-bottom plates (Corning, Wiesbaden, Germany) at a density of 10^4^cells/well and cells were maintained in cell-specific medium for 4 days. At day 4, cells were exposed to serum or CSF (v/v 25%) at different time-points as depicted in the figures. For analysis of P-gp up-take, cells were exposed to the P-gp specific substrate Rhodamine 123 [5µM] (Sigma-Aldrich, Taufkirchen, Germany) for 1 hr. As a positive control, a 30 min. pretreatment with the P-gp inhibitor PSC833 [0.5µM] (Sigma-Aldrich, Taufkirchen, Germany) was performed in all experiments and with every plate. After Rhodamine incubation, the cultures were washed three times with ice-cold PBS and cell-layer was then lysed in 200µL Triton-X100 (1% v/v in H_2_O). Data acquisition was performed with a fluoroluminometer (SpectraMax M2, Molecular Devices, San José, CA, USA) at following settings: λ absorbance at 485 nm, λ emission at 535 nm, cut-off > 530 nm, endpoint bottom read. Data processing was performed with the software SoftMax Pro 6 (Molecular Devices, San José, CA, USA). For the analysis of BCRP transporter function, the protocol was in principle similar to that of Rhodamine up-take measurement. As fluorescent substrate Mitoxantrone [20µM] (Sigma-Aldrich, Taufkirchen, Germany) was used. As positive control, the BCRP specific inhibitor KO143 [5µM] (Sigma-Aldrich, Taufkirchen, Germany) was used instead. Plates were measured with the following parameters: λ absorbance at 610 nm, λ emission at 685 nm, cut-off > 633 nm, endpoint bottom read.

### Activation of caspase activity in brain endothelial cells

The induction and differentiation of iPSC-derived brain endothelial cells (iPS-EC) has been described above. To indirectly analyse the induction of apoptosis in iPS-EC cells in vitro, caspase 3/7 activity (Caspase-Glo® 3/7 Assay Systems, Promega, Walldorf, Germany) was measured by cleaving a luminogenic substrate containing a DEVD amino acid sequence (Z-DEVD aminoluciferin). iPS-EC cells were induced, maintained and seeded in coated 96 flat-bottom plates (Corning, Wiesbaden, Germany) as described above. Cells were exposed to patient’s serum and CSF for 24 h as described above. Supernatants were removed and cells were washed three times with ice-cold PBS. The cell-layer was lysed and processed according to the manufacturer’s guidelines. Intrinsic Caspase 3/7 activity of serum and CSF probes was measured in parallel. Luminescence was detected using the fluoroluminometer (SpectraMax M2, Molecular Devices, San José, CA, USA) with the following settings: Lm1, 30 reads per well, PMT Gain “medium”. Data processing was performed with the software SoftMax Pro 6 (Molecular Devices, San José, CA, USA).

### Statistics and pathway analysis

Statistical analyses were performed using the SPSS® software package (SPSS version 21; SPSS Inc) as followed: Student’s *t*-test for quantitative normal data (QAlb; factor concentration by multi-bead array assay; all cell culture assays) and Mann–Whitney *U*-test for ordinal data (patient’s age; CSF cell number; mRS). Correlation analysis (spearman-rho) was performed for any significantly regulated factor with the grade of patient’s disability measured as mRS. Statistical significance was defined as two-tailed *P*-value of < 0.05. For further detailed description, see also **S1 Methods Appendix**.

To identify specific inflammatory networks in which the measured and differentially regulated factors are embedded, data were entered into the EMBiology^®^™ software (formerly Pathway Studio, Elsevier, New York). Bonferroni correction for multiple testing was applied in all cases. For further details on gene set enrichment analysis (GSEA), please refer to the **S1 Methods Appendix** [23, 24].

For subgroup analysis, patients in the infection cohort (VME and BME) were dichotomised into favourable (RS ≤ 4) and unfavourable (RS > 4) infection course. To exclude pathogen-specific effects on inflammatory factor levels in the dichotomised subgroups, only pairs of BME and VME with the same level of disability (mRS) were included. Using this approach, these defined subgroups were too small for GSEA. However, to identify associated pathways, we performed a structured Boolean search strategy in Pubmed (see **S1 Methods appendix** for detailed description).

In all figures, *p*-values are given as: * < 0.05, ** < 0.01, *** < 0.001 or depicted as absolute values. In all figures, factors are depicted according to HUGO Gene Nomenclature Committee (HGNC), the resource for approved human gene nomenclature. For the entire name of a given factor, we refer to http://www.genenames.org.

## Acknowledgments

We are grateful to Ludmila Umansky for her excellent technical assistance and to Tina Ohm for her continuous support as study nurse.

## Data availability statement

Most relevant data are within the paper and its Supporting Information files. In addition, multiplex (serum + CFS) and clinical raw data have been uploaded on DRYAD (https://datadryad.org/stash).

## Related manuscript statement

The authors state that they do not have a related or duplicate manuscript under consideration (or accepted) for publication elsewhere.

## Author contributions

**Conceptualization:** Thorsten Lenhard.

**Data curation:** Christine S. Falk, Viktor Balzer, Thorsten Lenhard.

**Formal analysis:** Thorsten Lenhard.

**Funding acquisition:** Uta Meyding-Lamadé, Thorsten Lenhard.

**Investigation:** Marie-Therese Herkel, Christine S. Falk, Viktor Balzer, Gert Fricker, Thorsten Lenhard.

**Methodology:** Christine S. Falk, Viktor Balzer, Gert Fricker, Thorsten Lenhard.

**Project administration:** Marie-Therese Herkel, Thorsten Lenhard, Corinna Schranz.

**Resources:** Christine S. Falk, Gert Fricker, Uta Meyding-Lamadé, Thorsten Lenhard.

**Supervision:** Christine S. Falk, Gert Fricker, Thorsten Lenhard.

**Validation:** Thorsten Lenhard, Christine S. Falk, Viktor Balzer

**Visualization:** Thorsten Lenhard

**Writing – original draft:** Thorsten Lenhard.

**Writing – review & editing:** Thorsten Lenhard, Marie-Therese Herkel, Christine S. Falk, Viktor Balzer, Gert Fricker, Corinna Schranz, Uta Meyding-Lamadé.

## Supporting information

**S1 Methods Appendix.** Detailed description of gene set enrichment analysis, Spearman-rho correlation analysis and structured Boolean search strategy.

**S1 Figure.** Correlation heatmap of baseline data and investigated factors in patients with VME and BME.

**S1 Table.** Overview of the cytokines, interleukins and growth factors that have been studied.

**S2 Table.** The raw data from the correlation analysis that served as the basis for the correlation heat map is shown in S1 Figure.

**S3 Table.** Host (patient) dependent signaling pathways and associated biological functions in patients with meningoencephalitis. – Boolean search strategy and reference list for Fig 4.

**S4 Table.** Impact of study results at the level of the neurovascular unit - Boolean search strategy and reference list for Fig 10.

